# A transcriptional constraint mechanism limits the homeostatic response to activity deprivation in mammalian neocortex

**DOI:** 10.1101/2021.10.20.465163

**Authors:** Vera Valakh, Derek Wise, Xiaoyue Aelita Zhu, Mingqi Sha, Jaidyn Fok, Stephen D. Van Hooser, Robin Schectman, Isabel Cepeda, Ryan Kirk, Sean O’Toole, Sacha B. Nelson

**Affiliations:** Department of Biology and Program in Neuroscience, Brandeis University, United States; Friedrich Miescher Institute for Biomedical Research, Switzerland

## Abstract

Healthy neuronal networks rely on homeostatic plasticity to maintain stable firing rates despite changing synaptic drive. These mechanisms, however, can themselves be destabilizing if activated inappropriately or excessively. For example, prolonged activity deprivation can lead to rebound hyperactivity and seizures. While many forms of homeostasis have been described, whether and how the magnitude of homeostatic plasticity is constrained remains unknown. Here we uncover negative regulation of cortical network homeostasis by the PARbZIP family of transcription factors. In cortical slice cultures made from knockout mice lacking all three of these factors, the network response to prolonged activity withdrawal measured with calcium imaging is much stronger, while baseline activity is unchanged. Whole cell recordings reveal an exaggerated increase in the frequency of miniature excitatory synaptic currents reflecting enhanced upregulation of recurrent excitatory synaptic transmission. Genetic analyses reveal that two of the factors, Hlf and Tef, are critical for constraining plasticity and for preventing life-threatening seizures. These data indicate that transcriptional activation is not only required for many forms of homeostatic plasticity but is also involved in restraint of the response to activity deprivation.

## Introduction

Neuronal networks are equipped with a set of homeostatic plasticity mechanisms that enable them to rebalance activity following perturbations during development, learning, or disease. Homeostatic plasticity can alter the intrinsic excitability of individual neurons, as well as the strength and number of both excitatory and inhibitory synapses (Davis, 2006; Turrigiano and Nelson, 2004). Synaptic changes can occur post-(Turrigiano et al., 1998) or presynaptically(Delvendahl and Müller, 2019) and may scale quantal amplitudes and alter the frequency of quantal events. These changes are homeostatic because they occur in the direction needed to rebalance the network after activity perturbation. Especially during development, homeostatic mechanisms are strong and can be maladaptive if they overshoot or are activated inappropriately (Nelson and Valakh, 2015).

Despite recognition of the significance of homeostatic plasticity, whether and how its strength is normally constrained has remained unknown. Downregulation of the strength of homeostatic plasticity could potentially provide protection against inappropriate or excessive activation of these mechanisms. However, such negative regulators of homeostatic plasticity have not been previously described.

Here we investigate the role of the proline and acidic amino acid-rich basic leucine zipper (PAR bZIP) family of transcription factors (TF) in homeostatic plasticity. The family consists of hepatic leukemia Factor (HLF), thyrotroph embryonic factor (TEF), and albumin D-site-binding protein (DBP) which are thought to act as transcriptional activators, as well as E4 Promoter-Binding Protein 4 (E4BP4, currently known as Nfil3) that acts as a transcriptional repressor (Mitsui et al., 2001). All 4 family members share the same DNA binding motif. DBP is a circadian gene controlled by CLOCK and oscillates with circadian rhythm (Ripperger et al., 2000). Other family members are also controlled by CLOCK (Li et al., 2017). However, while these TFs typically have oscillatory behavior in peripheral tissue, they do not oscillate in the brain outside of the SCN (Gachon et al., 2004). Loss of HLF, TEF and DBP has been associated with epilepsy (Gachon et al., 2004; Hawkins and Kearney, 2016; Rambousek et al., 2020) suggesting they are important in regulating network activity.

In order to investigate the role that gene transcription plays in regulating the neuronal response to activity deprivation, we profiled changes in gene expression engaged during homeostatic plasticity and found that prolonged activity deprivation activates a robust transcriptional program involving the PARbZIP TF family members HLF and TEF. Both act to restrain the expression of homeostatic plasticity. While they have limited effects on network function at baseline, they strongly suppress the upregulation of homeostatic changes, mainly by regulating recurrent excitatory synaptic connections, although inhibition is also affected. Together these results indicate that homeostatic plasticity is itself subject to activity-dependent regulation and is transcriptionally restrained by the PARbZIP TFs HLF and TEF.

## Results

### PARbZIP transcripts increase during activity deprivation

Certain forms of homeostatic plasticity require transcription (Goold and Nicoll, 2010; Ibata et al., 2008) and activity perturbation results in changes in gene expression (Schaukowitch et al., 2017). We hypothesized that among the differentially expressed genes, some induce or regulate homeostatic changes. To identify novel genes involved in the homeostatic response to activity deprivation, we measured gene expression in excitatory and inhibitory neurons in organotypic slice cultures following a global decrease in activity. We cut coronal slices including neocortex at P7 and cultured them for 5 days. During this culture period the neurons reform connections, and spontaneous bouts of activity, termed up states, emerge (Johnson and Buonomano, 2007; Koch et al., 2010). To broadly block activity, we applied the voltage-gated sodium channel blocker tetrodotoxin, TTX (0.5 μM), for 5 days during an early developmental period (equivalent postnatal day, EP, 12-17). This manipulation induces a robust homeostatic program and profoundly changes network dynamics (Koch et al., 2010; Schaukowitch et al., 2017). To characterize the changes in gene expression, we sorted deep layer (L5 and L6) pyramidal and PV+ interneurons and performed RNA sequencing to look for transcripts that change following prolonged silencing. Inactivity activates a robust transcriptional program of both positively and negatively affected genes (Figure 1A) consistent with data from dissociated hippocampal cultures (Schaukowitch et al., 2017). To identify transcriptional regulators of homeostatic plasticity, we looked for transcription factors (TFs) upregulated in the TTX condition. TFs can be classified into families, and in some cases subfamilies, which share a DNA binding domain, and so are expected to bind the same target DNA sequences in the genome and therefore to regulate the same or similar target transcripts. We asked which families or subfamilies were most over-represented amongst the TFs differentially expressed during activity blockade. The most overrepresented subfamily or family was the PARbZIP subfamily of TFs, a subtype of the CEBP-related family in the class of Basic leucine zipper factors (classification data from tfclass.bioinf.med. uni-goettingen.de (Wingender et al., 2015). This subfamily includes 3 transcriptional activators Hlf, Tef, and Dbp, and a transcriptional repressor, Nfil3. Expression levels of Dbp are much lower than those of Hlf and Tef (<3 TPM in excitatory and PV+ inhibitory neurons), suggesting that this family member may play a more limited role in the neocortex. Both Hlf and Tef are robustly upregulated after 5 days of activity deprivation in both excitatory (Fold Change TTX/control; FC = 2.6, 2.5; adjusted p-value; padj = 1.3e-8, 3.8e-4,) and inhibitory neurons (FC = 7.0, 2.1; padj = 4.6e-39, 2.0e-3), while Nfil3 and Dbp were not (FC = 0.51,1.9; padj= 0.49, 0.80, excitatory; FC= 1.2, 1.4; padj= 0.84, 0.79 inhibitory).

**Figure 1.**
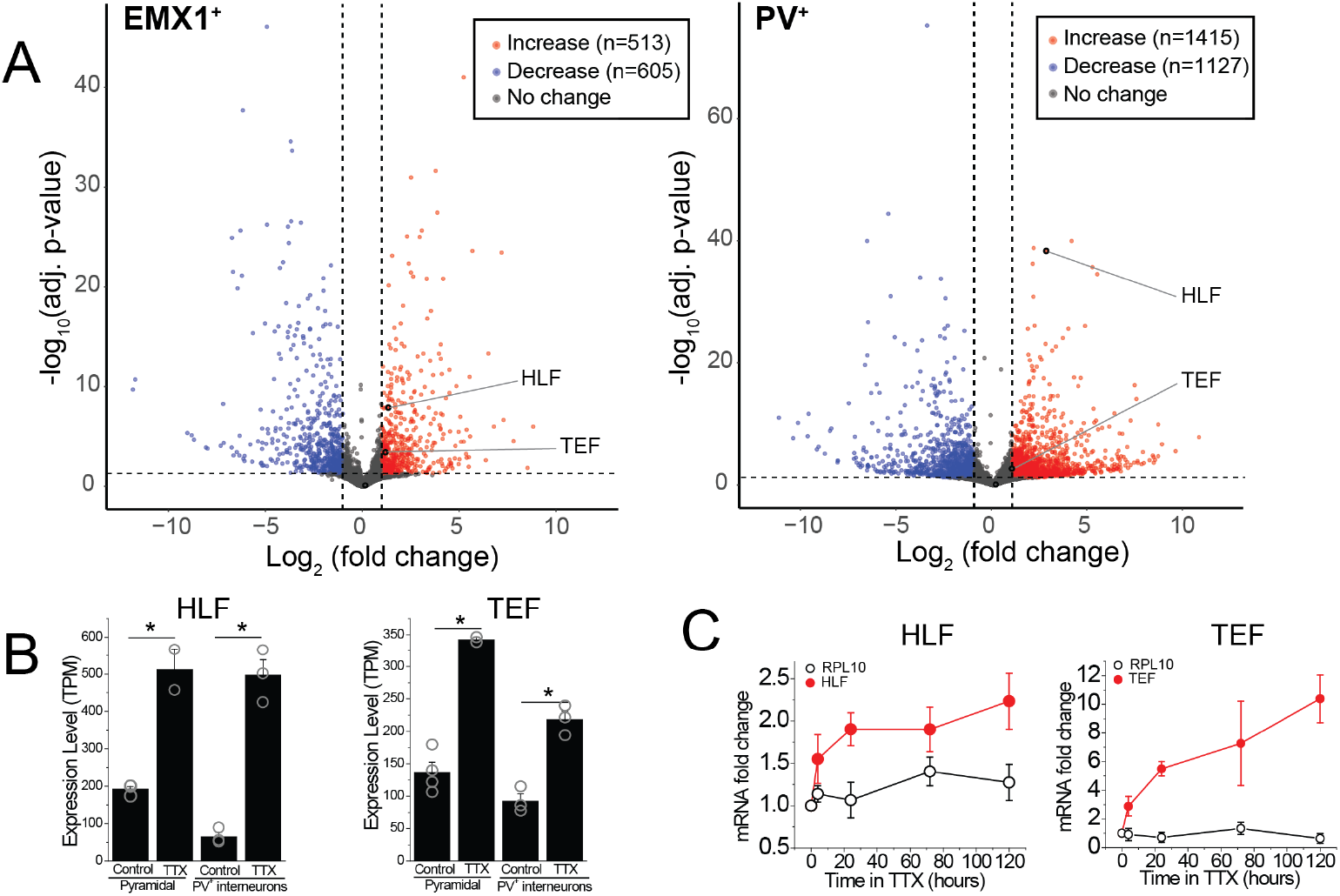
RNA-sequencing identifies transcripts affected by activity deprivation including the PARbZIP family of transcription factors. (A) Volcano plots of bulk RNA-seq from sorted fluorescently labeled pyramidal cells (left) and PV+ interneurons (right). Dashed lines indicate a Fold-change of 2 and adjusted p-value of 0.05. Differential expression analysis revealed that TTX resulted in upregulation (513 in pyramidal, 1415 in PV+ interneurons) as well as downregulation (605 in pyramidal, 1127 in PV+ interneurons) of genes, among which Hlf and Tef were identified as upregulated with activity block in both excitatory and inhibitory neurons. Statistical analysis (Wald test followed by Benjamini-Hochberg correction) reveals that Hlf and Tef are significantly upregulated in both pyramidal cells and PV+ interneurons, while Dbp and Nfil3 are not significantly altered in either cell type. (B) Bar graphs displaying transcript per million (TPM) values of Hlf (left panel) and Tef (right panel) in pyramidal cells (left) and interneurons (right) in control and 5-day TTX treated slices. Bars are mean values, open symbols are individual experiments. (C). Quantitative real-time PCR analysis of Hlf and Tef expression in whole cortex lysates following a time course of activity deprivation (n=3-4 slices per time point). Two-way ANOVA reveals significant differences between Hlf/Tef and RPL10 (p<0.05).

To validate this finding and to further dissect the time course of PARbZIP transcription factors expression, we isolated the whole cortex from slice cultures and measured Hlf and Tef transcript levels at various time points during activity deprivation using real-time quantitative PCR. We find that upregulation occurs soon after activity withdrawal, since by 4 hours of TTX application the level of both TFs is already substantially elevated (Figure 1C), consistent with previous findings in dissociated hippocampal cultures (Schaukowitch et al., 2017). We also find that Hlf and Tef continue to increase with time, suggesting that their upregulation correlates with how long the network has been silenced.

### PARbZIP TFs restrain network homeostatic plasticity

Since Hlf and Tef are progressively upregulated during prolonged TTX treatment, which can lead to subsequent epilepsy (Galvan et al., 2000; Scharfman, 2002), we entertained the hypothesis that they may contribute to the homeostatic increase in network excitability following activity deprivation. On the other hand, loss of function of members of the PARbZIP family of TFs are also associated with epileptic phenotypes (Hawkins and Kearney, 2016, 2012; Rambousek et al., 2020). Specifically, triple knockout (TKO) Hlf-/-/Dbp-/-/Tef-/- mice have epilepsy (Gachon et al., 2004). One possibility that reconciles this apparent conflict is that instead of driving homeostatic plasticity, Hlf and Tef are upregulated in the TTX condition to restrain homeostatic responses. To test this hypothesis, we investigated the role that these TFs play in homeostatic plasticity. We induced homeostatic upregulation of network function by two-day TTX application in organotypic slice cultures and measured network activity after TTX removal.

As a read-out of network activity, we measured changes in intracellular calcium using virally-delivered GCaMP6f in primary somatosensory cortex (Figure 2A). In untreated slices at baseline, the neurons display population activity organized into complex, infrequent up states which are similar to those observed in vivo (Figure 2B, Sanchez-Vives and McCormick, 2000). To quantify network activity, we measured the frequency of Ca2+ peaks in each recording and the synchrony of calcium transients across cells (see methods for detailed description of quantitative analysis). After two days of TTX treatment, the frequency of the up states increases. While 2 days of TTX incubation increases the peak frequency 3.6-fold in WT slices (from 0.06 +/- 0.01 to 0.22 +/- 0.04 Hz), the same silencing duration produces a nearly six-fold increase in the TKO slices (from 0.09 +/- 0.02 Hz to 0.5 +/- 0.05 Hz; Figure 2, Figure 2—Figure Supplement 1A). A two-way ANOVA revealed significant effects of treatment (TTX vs. Control; p<0.0001) and genotype (p<0.0001) and a significant interaction between the two (p<0.0001). Post hoc T-tests (with Tukey correction for multiple comparisons) revealed that TTX increases in activity were highly significant in both WT (p= 0.0007) and TKO (p<0.0001) slices and that TTX produced a significantly stronger increase in the TKO (p<0.0001), while the baseline activity of cells from slices derived from the TKO and WT do not differ significantly (p=0.92). Consistent with this, normalizing the TTX response to the control response revealed a larger relative increase in TKO than WT cultures (Fig. 2—Figure supplement 1A). The TTX condition also increased synchrony (from 0.76 +/- 0.03 to 0.95 +/- 0.01) in WT, and in the TKO (from 0.74 +/- 0.03 to 0.94 +/- 0.05; Figure 2—Figure Supplement 1B). While the TTX effects were highly significant (Two-way ANOVA; p<0.0001), there was no significant effect of genotype (p=0.43) and no interaction between treatment and genotype (p=0.65), presumably reflecting the fact that synchrony is already saturated by TTX treatment in the WT and cannot further increase in the TKO, but also indicating that baseline synchrony is not altered in the TKO. A power test revealed that with the effect size and variances observed, 95 slice cultures in each group would be needed to detect a significant difference between WT and TKO peak frequency, while for the synchrony measure, the required N would be 1609. These data suggest that the PARbZIP family of TFs normally functions to restrain homeostatic plasticity, and in their absence, the homeostatic response is exaggerated. Moreover, even though the mutation is associated with epilepsy, it does not make cortical neuronal circuits more excitable at baseline.

**Figure 2.**
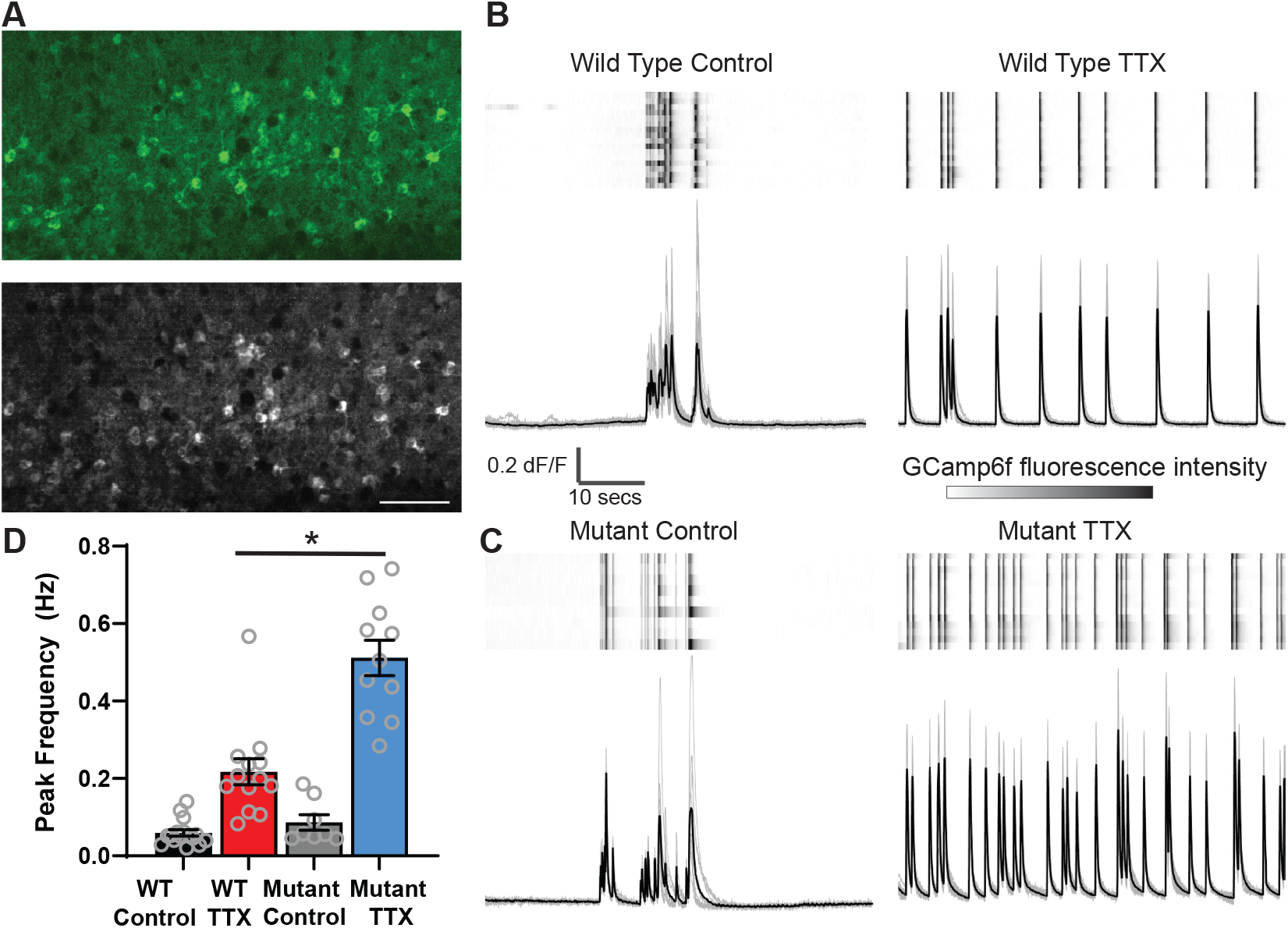
Calcium imaging reveals exaggerated response to activity deprivation in Hlf-/-/Dbp-/-/Tef-/- triple knockout (TKO) slices. (A). A confocal image of GCaMP6f fluorescence during an up state in wild type (WT) control slice (top panel) and a standard deviation projection of the same field of view of calcium signal during one minute of recording (bottom panel). Scale bar =50 μm. ROIs were manually selected around active cells identified by high standard deviation values. (B,C) GCaMP6f fluorescence heat map of selected ROIs (rows) during one minute of recording (top panels) also shown as overlapping traces (gray, bottom panels). and an average fluorescence (black trace, bottom panels) during one minute of recording in WT (B) and TKO (mutant) slices (C) from control (left) and TTX-treated conditions (right). (D) Quantification of peak frequency in GCaMP6f fluorescence traces. N = 16 slices for WT Control, N = 13 WTT TX, N = 8TKO Control, N = 11 TKO TTX. TTX treatment significantly increases peak frequency in WT slices with significant (p<0.0001) interaction between treatment and genotype. The increase is more dramatic in the TKO slices. Two-way ANOVA with Tukey’s correction for multiple comparisons to test statistical significance between conditions * p≤0.0001. **Figure 2—figure supplement 1.** Ca2+ activity in mutant slices is similarly exaggerated after 2 and 5 days of activity withdrawal. **Figure 2—figure supplement 2.** Ca2+ activity recovers to baseline levels following washout of TTX in both wild type and mutant slices.

### Recovery from activity deprivation is unperturbed in the mutant slices

Homeostatic plasticity is thought to provide flexibility to the network and thus most forms are reversed after reintroduction of activity (Desai et al., 2002; Hobbiss et al., 2018; Koch et al., 2010; Wallace and Bear, 2004). Reversal involves a homeostatic reduction of abnormally elevated activity. Even when this involves symmetric opposing changes in the same biophysical parameters altered in the response to activity deprivation (such as with downscaling and upscaling of excitatory quantal amplitudes), the mechanisms involved typically differ (Stellwagen and Malenka, 2006; Sun and Turrigiano, 2011; Tan et al., 2015; Wang et al., 2017). In addition, changes in activity following rebound from deprivation could reflect either a change in the vigor of the circuit’s attempt to restore activity (i.e. enhanced homeostatic plasticity) or a persistent change in the setpoint to which activity is returned (Styr et al., 2019), or both. To test whether the transcriptional restraint of homeostasis is required for bidirectional flexibility and to assess whether the setpoint had changed, we asked whether the network is able to return back to baseline activity levels when the activity blockade is removed. We measured network activity immediately after the end of silencing with TTX as well as following two days of recovery in TTX-free media. In WT slices, activity deprivation causes hyperactivity, evident from a two-fold rise in the peak frequency of the calcium activity. However, this exuberant activity is restored back to baseline levels within two days following restoration of action potential firing. In the TKO slices, even though the initial response to TTX is exaggerated, the network is also able to return to baseline levels following two days of recovery, to levels indistinguishable from WT (Figure 2—Figure Supplement 2). These data suggest that the recovery from a high activity state is not diminished in the TKO and suggests that the mutation in fact produces exaggerated homeostatic plasticity rather than a change in activity setpoint.

### Frequency, but not amplitude, of mEPSCs is disproportionately upregulated by deprivation in TKO slices

Network activity is unaltered in the mutant at baseline but the response to activity deprivation is exaggerated. To learn more about the mechanism by which Hlf and Tef restrain homeostatic plasticity, we probed the effects of TTX on excitatory synaptic transmission. Because of the high levels of spontaneous network activity, it was not feasible to study unitary action potential evoked transmission. Instead, to broadly assay the properties of excitatory synapses, we measured pharmacologically isolated AMPA-receptor driven miniature EPSCs in control and TTX-treated slices. We observed a robust upregulation of mEPSC amplitude following two days of activity deprivation, consistent with our understanding of homeostatic synaptic scaling (Turrigiano et al., 1998). We also observed a large increase in mEPSC frequency, which has also been described in this preparation (Koch et al., 2010) and in multiple other preparations (Wierenga et al., 2006). In the TKO mice, the synaptic scaling is unaffected (Figure 3 A,C; amplitude increased from, 13.4 +/- 1.1 to 18.5 +/- 1.2 pA in WT and from 13.2 +/-1.4 to 19.7 pA +/-1.1 in TKO) but the increase in mEPSC frequency in response to activity deprivation is more dramatic (Figure 3 B,D 4.0 +/- 0.8 to 6.2 +/- 0.7 Hz in WT vs. 3.5 +/- 0.5 to 9.3 +/-1.0 Hz in the KO). Notably both the frequency and amplitude of mEPSCs are the same between WT and mutant slices at baseline which parallels our observations of network activity (Figure 2). These results suggest that the members of the PARbZIP family of TFs function to restrain upregulation of mEPSC frequency in response to activity deprivation and this increase is correlated with changes in activity levels across the network.

**Figure 3.**
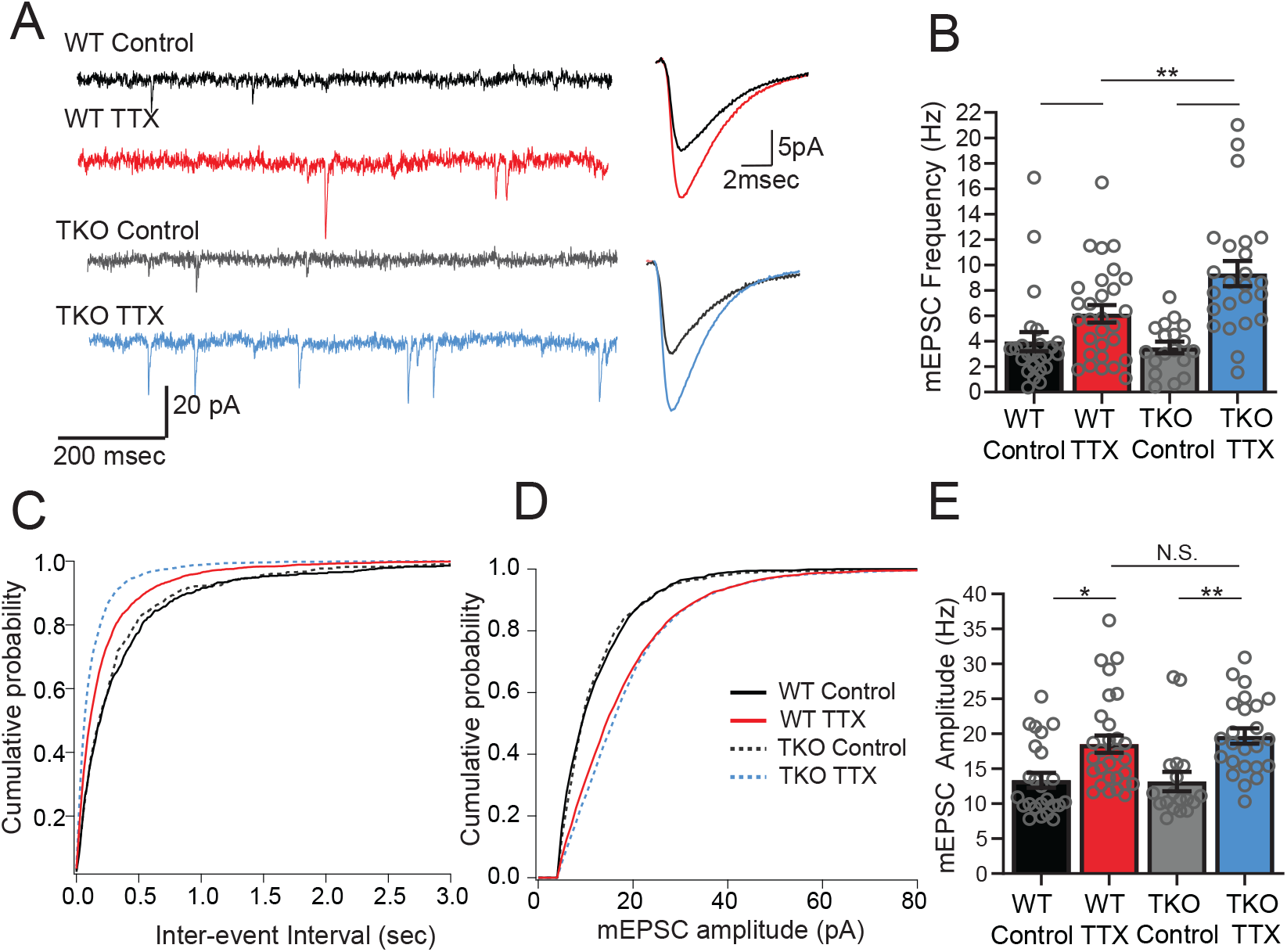
Frequency but not amplitude of excitatory synaptic currents is disproportionally upregulated in TTX-treated TKO slices. (A) Representative traces of mEPSC recordings from WT control (black), WT TTX-treated (red), TKO control (gray), and TKO TTX-treated (blue) slices. Right panel, average mEPSC waveforms for the same conditions. Both the frequency and the amplitude is increased in TTX-treated slices, however, the increase in frequency is more dramatic in TKO slices. (B) Quantification of mEPSC frequency for each condition. Colored bars with error bars are mean ± SEM, open circles are individual cellsN=24 for WT control, N=29 for WT TTX, N=18 for TKO control, N=24 for TKO TTX. Two-way ANOVA revealed significant main effect of TTX treatment (p<0.0001) and interaction between treatment and genotype (p=0.023) but not the genotype alone (p=0.089). Post hoc Tukey test revealed enhanced effect of TTX on mEPSC frequency in TKO cells compared to WT (p=0.016, *).(C) Cumulative probability histogram for mEPSC inter-event intervals in each condition. (D) Cumulative probability histogram for mEPSC amplitudes in each condition. (E) mEPSC amplitudes for each condition. Two-way ANOVA revealed significant main effect for drug treatment (p<0.0001) but not genotype (p=0.69) or interaction between treatment and genotype (p=0.59). TTX treatment enhanced mEPSC amplitude in both WT (p=0.01, Tukey post hoc test, *) and TKO (p=0.0033, Tukey post hoc test, **) to the same extent (p=0.89, WTTTX compared to TKO TTX, Tukey post hoc test, N.S.)

### Neither the frequency nor amplitude of mIPSCs in pyramidal neurons in the TTX condition are affected in the TKO

Since L5 pyramidal neurons also receive inhibitory input, which is also subject to homeostatic plasticity (Kilman et al., 2002; Kim and Alger, 2010), we wanted to determine whether the response of mIPSCs to activity deprivation is also exaggerated in the mutant mice. In wild-type cultures, measurement of inhibitory input onto L5 pyramidal neurons revealed that both the amplitude and frequency of mIPSCs drop in response to activity deprivation (Figure 4). When we measured mIPSCs in pyramidal neurons from TKO slices, we saw a similar decrease in frequency following 48 hr TTX treatment, although the amplitude of the change was smaller than in the WT (Figure 4 B,C). In contrast, the mIPSC amplitude change is abolished: mIPSC amplitude is decreased at baseline compared to WT but did not show a further decrease after TTX treatment (Figure 4 D,E). However, the decreased amplitude in the untreated condition does not correlate with the unchanged baseline network activity levels (Figure 2), suggesting that the decrease of mIPSCs amplitude at baseline may be insufficient on its own to cause network hyperexcitability. Thus, both the amplitude and frequency of mIPSCs received by pyramidal neurons after activity deprivation are indistinguishable from the wild-type in the TKO slices, suggesting that homeostatic plasticity of mIPSCs is not directly driving the changes in the network activity following TTX treatment. We cannot rule out the possibility that although baseline activity is normal, the baseline reduction in mIPSC amplitude contributes to subsequent network plasticity indirectly, by contributing to induction of the changes in EPSCs observed, or by inducing some form of compensation other than the physiological parameters measured.

**Figure 4.**
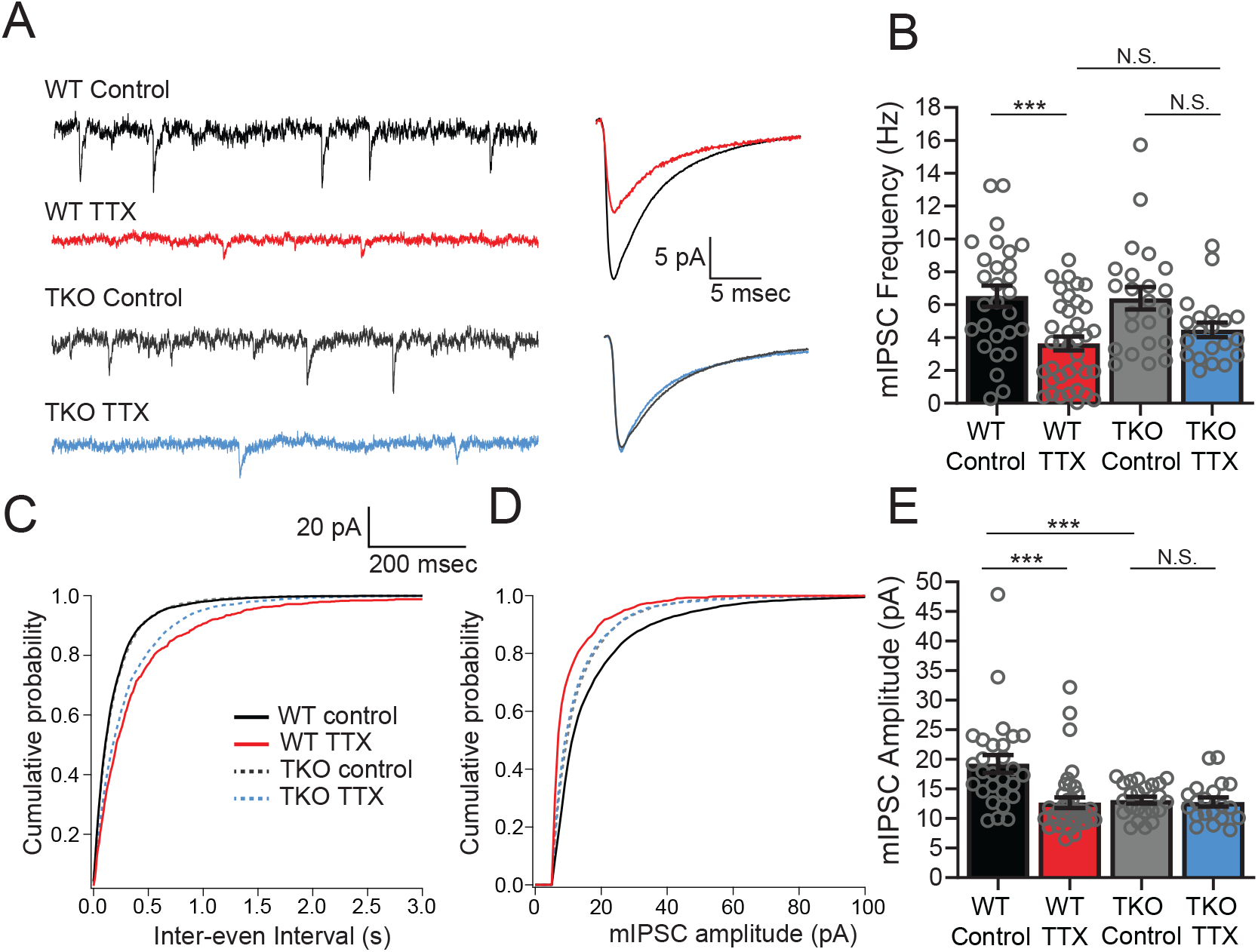
Inhibitory synaptic currents are not affected by the Hlf-/-/Dbp-/-/Tef-/- mutation in the TTX condition. (A) Representative traces of mIPSC recordings from WT control (black), WT TTX-treated (red), TKO control (gray), and TKO TTX-treated (blue) slices. Right panel, average mIPSC waveforms for the same conditions. TTX treatment results in a diminished effect on mIPSC frequency in TKO and does not further decrease the amplitude of mIPSCs in TKO slices. (B) Quantification of mIPSC frequency. Colored bars with error bars are mean ± SEM, open circles are individual cells. Two-way ANOVA revealed a significant main effect of TTX treatment (p<0.0001) but not genotype (p=0.54). There was no significant interaction between treatment and genotype (p=0.40), N=28 for WT control, N=37 for WT TTX, N=23 for TKO control, N=20 for TKO TTX. TTX treatment decreased mIPSC frequency in WT (p=0.0007, post hoc Tukey test) but not in TKO cells (p=0.14, post hoc Tukey test). mIPSC frequency in TTX-treated cells is not different in TKO slices compared to WT (p=0.72). The normalized change in frequency (TTX normalized to Control) was 0.55 in WT and 0.70 in TKO. We performed a power analysis that revealed that for the observed effect size and variance a sample of 68 neurons per condition would be required for significance. In B and E, (C) Cumulative probability histogram for mIPSC inter-event intervals in each condition. (D) Cumulative probability histogram for mIPSC amplitudes in each condition. (E) Quantification of mIPSC amplitudes for the same conditions as in B. Two-way ANOVA revealed significant main effect of TTX (p=0.002), and genotype (p=0.0067), and a significant interaction between treatment and genotype (p=0.005). TTX treatment decreased mIPSC amplitude in WT (p<0.0001, Tukey post hoc test, ***), but not in TKO cells (p=0.997, Tukey post hoc test, N.S.) presumably due to significantly lower baseline mIPSC amplitude in TKO slices compared to WT (p=0008, post hoc Tukey test, ***).

### Intrinsic excitability of pyramidal neurons is unaffected in TKO slices

Changes in neuronal excitability have also been described as part of the homeostatic response to activity deprivation (Desai et al., 1999; Karmarkar and Buonomano, 2006; Lambo and Turrigiano, 2013; Moore et al., 2018; Yoshimura and Rasband, 2014). We wanted to test whether changes in intrinsic excitability could be contributing to the exaggerated homeostatic response in the TKO slices similarly to mEPSCs. WT L5 pyramidal neurons become more excitable following prolonged activity deprivation (Figure 5; Desai et al., 1999). To account for difference in the proportion of adapting and non-adapting neurons (Hattox and Nelson, 2007) causing apparent changes in intrinsic excitability due to uneven sampling of the two groups, we measured the adaptation ratio of each cell and analyzed an equal number of adapting and non-adapting cells in each genotype. Changes in excitability due to activity deprivation are accompanied by a shift in the average adaptation ratio of the cells in both WT and TKO slices (control WT: 3.8±0.16, TTX WT: 0.72 ±0.22, TKO control: 0.46±0.25, TKOTTX: 0.71 ±0.33, two-way ANOVA significant difference between control and TTX and no significant difference between WT and TKO). Excitatory neurons in L5 in the slices prepared from TKO animals are equally excitable at baseline and also show upregulation of intrinsic excitability to the same extent (Figure 5). These results suggest that changes in intrinsic excitability are not driving the exaggerated network response to activity deprivation and this form of homeostatic plasticity is not under the control of the Par bZIP TF family.

**Figure 5.**
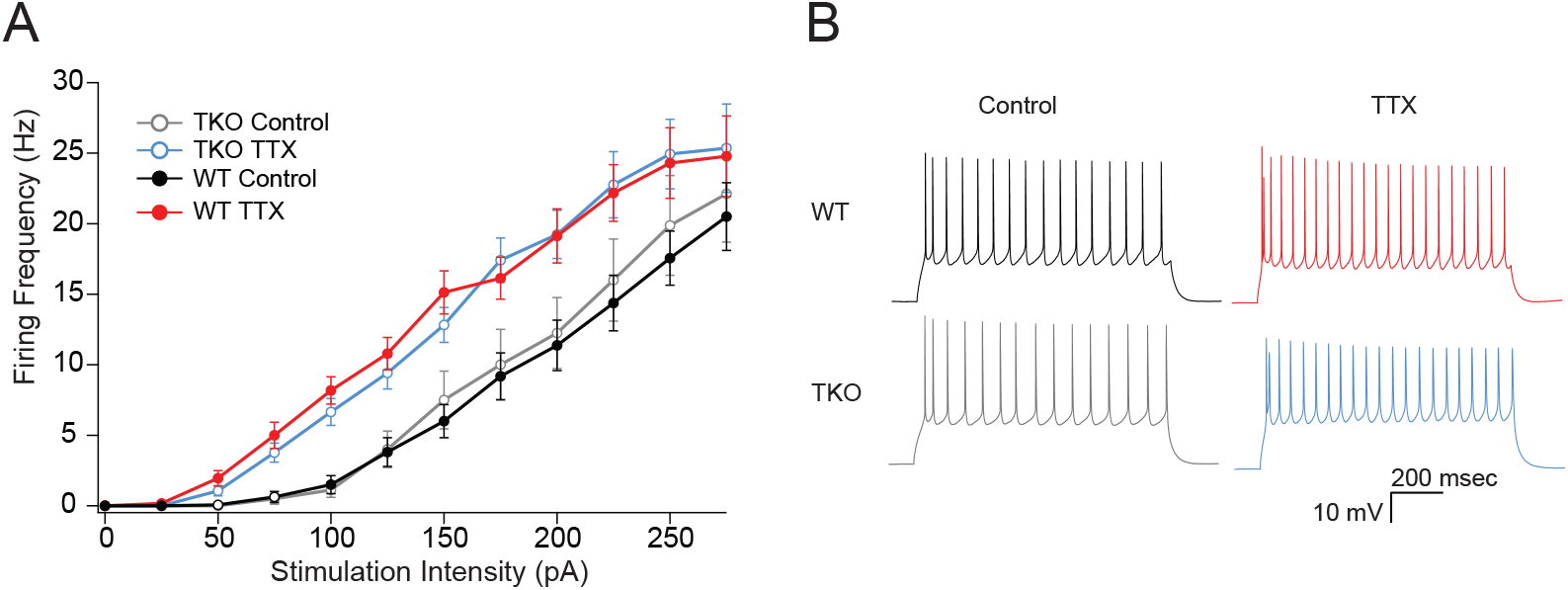
The effects of TTX on intrinsic excitability are not altered in TKO pyramidal neurons. (A) Comparisons of frequency-current relationships for layer 5 pyramidal neurons in control (black, closed circles) and TTX-treated (red, closed circles) WT slices, and in control (gray, open circles), and TTX-treated (blue, open circles) TKO slices, N=27 cells for WT TTX, N= 16 for WT control, N=8 for TKO control, N=17 for TKO TTX. Two-day TTX treatment increases intrinsic excitability in both WT and TKO slice cultures to a similar extent. A 3-way mixed ANOVA with between subjects factors of treatment and genotype, and a within subjects factor of current level, revealed significant main effects for TTX treatment (F(1,60) = 18.1, p =7.3e-05) and current level (F(11,660) = 2.3e-190) but not for genotype (F(1,60) = 0.06, p=0.94. There was a significant interaction between treatment and current level (p=.06) but not for interactions of genotype with current level (p=0.93) or with both current level and treatment (p=0.99). (B) Example traces of a train of action potentials from a pyramidal neuron in response to a 175pA 0.5s depolarizing current injection in control (left), TTX-treated (right), WT (top), or TKO (bottom) slices.

### PARbZIP family members play unequal roles in restraining homeostatic plasticity

Expression of both Hlf and Tef increase in excitatory neurons following activity deprivation. Because all three Par bZIP family members form homo- and heterodimers and bind to the same PAR-response element (Gachon, 2007), they may be able to compensate for one another. To test whether Hlf, Tef, and Dbp work together to regulate homeostatic plasticity or if they have unequal contributions to restraining network response to activity deprivation, we measured the homeostatic response while varying gene copy number of each transcription factor. In addition to the WT and TKO data described in Fig 2 and Fig 2—Figure Supplement 1, we measured calcium transients in slice cultures treated with or without TTX from 97 additional animals with various combinations of 0, 1 or 2 alleles of Hlf, Tef and Dbp (full details of each experiment given in associated data file). Rather than attempt to statistically test individual differences between each of 27 potential genotypes (Null, Het, wild type for the three genes considered jointly), we fit a series of multivariate linear models to test the ability of genotype to predict calcium peak frequency. The same six models, described in Table 1, were fit separately to data from control and TTX-treated slices. The models, shown in Table 1, differ in how they depend on the number of alleles of each of the three genes. Model 1 depends on all three genes, models 2-4 depend only on one of the three genes (each in turn) and models 5 and 6 depend only on Hlf and Tef, either independently, or summed together. The ability of each model to account for the data was estimated from the adjusted R2 which reflects the fraction of the variance in the data predicted by the model variables. Several key points were evident from this analysis:

1. All of the models produced poor fits (adjusted ***R***^2^ of −0.01 to 0.063) to the control data, consistent with the hypothesis that baseline activity does not depend on genotype.
2. All of the models that included variables for Hlf and/or Tef produced much better fits to the TTX data (adjusted ***R***^2^ of 0.198 to 0.296) consistent with the hypothesis that rebound activity is influenced by genotype. In these models, the coefficients (effect size) for Hlf and Tef were similar and T-tests revealed they were highly significantly different from zero.
3. The model that only depended on Dbp (line 4) provided a poor fit to the TTX data (adjusted ***R***^2^ = 0.073) and in the model that included both terms for Dbp and the other two factors (line 1), the Dbp coefficient was small and not significantly different from zero. These observations are consistent with the hypothesis that rebound activity does not depend on Dbp and that even in a significant model including Hlf and Tef, Dbp does not add additional explanatory power to the model.
4. The models that included terms for both Hlf and Tef (line 5) or for their sum (line 6) produced the best fits (adjusted ***R***^2^ = 0.291 and 0.296); i.e. better than those that depend on only one of these factors (lines 2,3; adjusted ***R***^2^ = 0.198, 0.229). This is consistent with the hypothesis that both genes contribute to constraining cortical homeostatic plasticity.

**Table 1.**
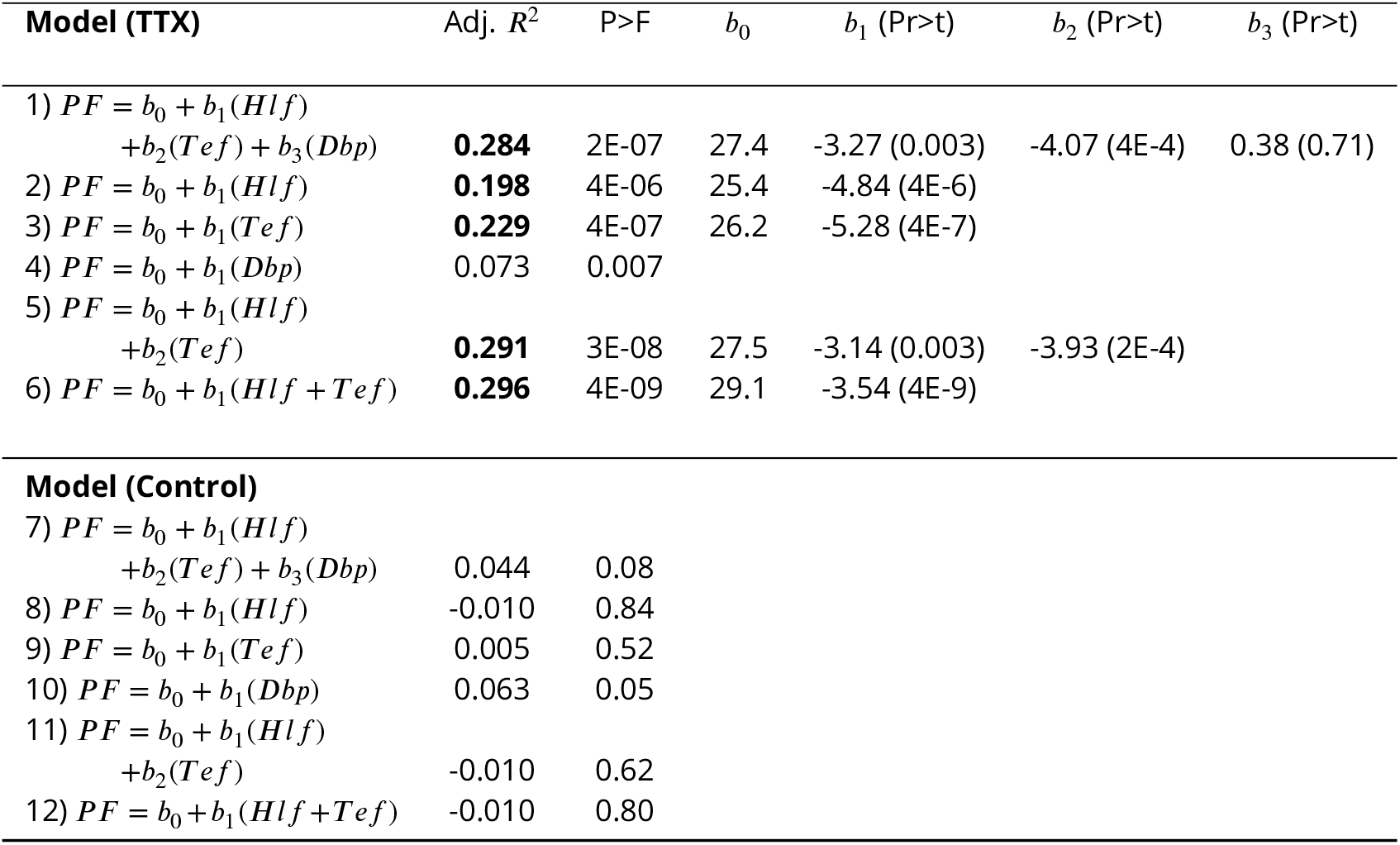
Multivariate linear models of the dependence of calcium peak frequency (PF) on genotype. Listed linear models were fit to peak frequencies of calcium transients measured from 87 control and 98 TTX-treated slices each from separate animals that differed in their genotype (null, heterozygote or wild type) for each of the *Hlf, Tef* and *Dbp* genes. The data (see attached file) includes the 88 experiments shown in Figure 2 and figure 2—Figure supp1ement 1 (WT and TKO) as well as 97 additional experiments (intermediate genotypes). Regressions were performed using the lm function in R where the variables Hlf, Tef and Dbp represent the numbers of alleles of each gene (0, 1 or 2) and the variable (Hlf + Tef) represented the sum of the number of Hlf and Tef alleles. The column p>F reports the significance of each model assessed from an F test. For models that accounted for > 10% of the variance in the data (Adj *R^2^* > 0.1; bold values), the intercept (*b*_0_) and coefficients (*b*_1_…) are listed, along with the probability that the coefficient is nonzero (T-test). Pr>t was <2E-16 for all *b*_0_ given.

Since the model with the highest adjusted ***R***^2^ was that which depended only on the sum of the number of Hlf and Tef alleles (line 6), we used this model for further post hoc (Tukey) tests to determine the effect of different numbers of alleles on the TTX response. The results of this analysis are shown in Figure 6. The most significant differences were between TKO slices (0 alleles; n=32, mean=0.48) and the other genotypes tested (1 allele, n=16, mean=0.32; 2 alleles, n=26, mean=0.35; and 4 alleles, n=24, mean=0.23; p <0.0001 to 0.0014). Most of the other comparisons were not significant, with the exception of the difference between WT responses and those for animals with only two alleles (p=0.011). This indicates that most of the difference between TKO (zero alleles of Hlf + Tef) and WT (4 alleles of Hlf + Tef) could be restored by a single allele, although there was a small but significant improvement between 2 and 4 alleles. The fact that this difference was significant, but the differences between 1 and 2 alleles, and 1 and 4 alleles were not, may reflect a real difference between these groups, or may reflect the imbalanced numbers of animals in each group. The group with 2 alleles included animals that were WT for Hlf but lacked Tef (N=8), animals that were WT for Tef and lacked Hlf (N=9), and trans-Het animals that were heterozygous for both Hlf and Tef (N=7). A separate analysis of variance of these three subgroups revealed that the means (0.32, 0.41 and 0.33) were not significantly different (p=0.28), consistent with the hypothesis that alleles of Hlf and Tef can substitute for one another in their ability to regulate the homeostatic response to activity deprivation.

**Figure 6.**
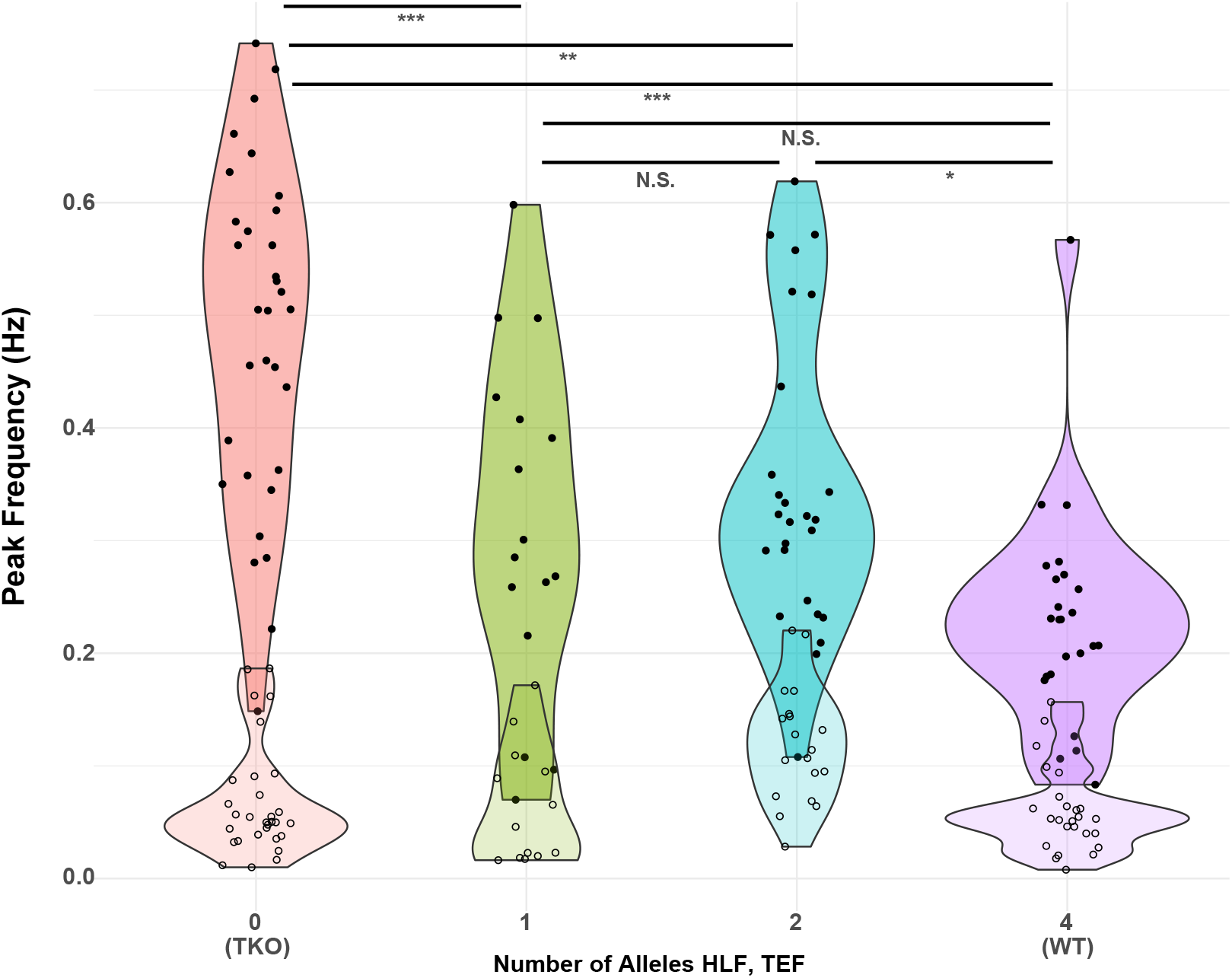
Figure 6. Presence of one allele of Hlf or Tef largely restores the WT response to activity deprivation. Violin plots show the distribution of peak frequencies measured from TTX-treated (upper, darker colors, filled symbols) and control (lower, lighter colors, open symbols) slice cultures made from animals carrying 0, 1, 2 or 4 alleles of the PARbZIP TFs Hlf and Tef. Data are those used in Table 1 with the X-axis corresponding to models 6 (TTX) and 12 (Control). Horizontal positions of individual data points are jittered to prevent overplotting. Horizontal lines at top indicate results of post hoc T-tests between levels for the TTX data with Tukey correction for multiple comparisons. *adj. p < 0.05, **adj. p < 0.01, ***adj. p < 0.001, N.S. – not significant, adj. p >0.05. No post hoc testing was performed for the Control data since the fit of model 12 was not significant.

To further dissect the relative contribution of the Par bZIP family members we measured the survival of the mice with different numbers of copies of the TFs. TKO animals develop spontaneous seizures and have a dramatically decreased lifespan (Gachon et al., 2004). While our objective was not to make a detailed study of the seizure phenotype, we also observed, on multiple occasions, TKO animals that experienced spontaneous seizures, even though for most of cases, the instance of death was not directly observed. However, just one allele of either Hlf or Tef restores the lifespan of the animals to levels not statistically different from WT. Mice with one Dbp allele have improved survival compared to TKO animals but still die at a statistically significant higher rate than WT mice (Figure 7). These results differ from the cortical network activity response to activity deprivation in organotypic slices where the presence of Dbp had no effect on the constraint of homeostatic plasticity. This suggests that even though Dbp is dispensable for restraining homeostatic plasticity in the cortex, it may play a role in preventing premature death by functioning elsewhere in the brain or peripheral tissue (Stewart et al., 2020) since these TF are expressed and play important roles there (Wahlestedt et al., 2017; Wang et al., 2010). However, a single allele of Hlf and Tef is enough to compensate for loss of the rest of the family members and restore survival to WT levels.

**Figure 7.**
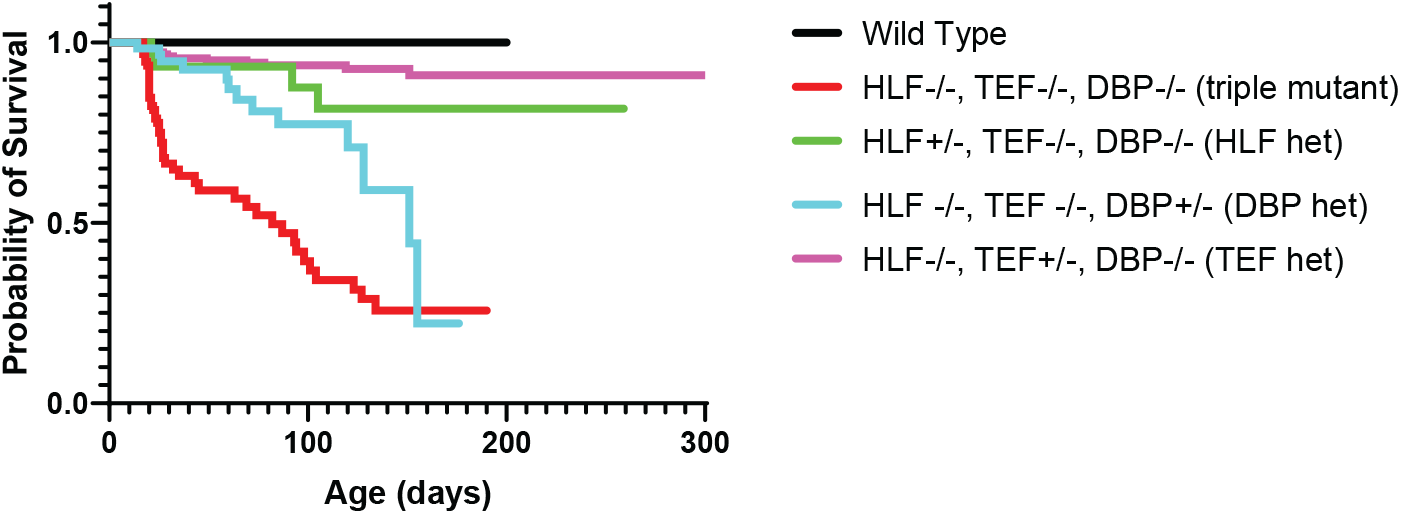
Presence of one allele of a PARbZIP TF improves survival relative to TKO animals. TKO mice have dramatically decreased survival (red line) compared to wild-type mice (black line). Presence of one copy of either Hlf (green line) or Tef (purple line) prevents premature death and the survival curves are not statistically different from wild-type, p = 0.12 and p = 0.56 respectively, Log-Rank test with Bonferroni correction for multiple comparisons. DBP heterozygous mice (blue line) also have improved survival over TKO mice (p = 0.003) but are still statistically more likely to die prematurely than wild-type mice (p = 0.0003). N=59 for DBP heterozygous, N=96 for TKO, N=36 for HLF heterozygous, N=23 for WT, N=179 for TEF heterozygous.

## Discussion

Inappropriately triggered homeostatic plasticity can either fail to compensate for changes in activity or can itself destabilize network activity (Nelson and Valakh, 2015). Although some molecular pathways that are required for the induction of homeostatic plasticity have been identified, whether and how homeostatic plasticity is negatively regulated is unknown. Here we show that reduced activity, which initiates a compensatory increase in network excitability, also activates a set of transcription factors that function to restrain homeostasis. Thus, we propose a model in which homeostatic plasticity is negatively regulated through the PARbZIP family of TFs.

Excitatory neurons respond to a global drop in network activity by upregulating the frequency and amplitude of excitatory synaptic currents, decreasing inhibitory currents and increasing intrinsic excitability. These changes correlate with dramatically increased network activity immediately following removal of activity blockade that “overshoots” initial activity levels. Synaptic scaling of excitatory synapses has previously been shown to depend on intact transcription and translation (Dörrbaum et al., 2020; Goold and Nicoll, 2010; Ibata et al., 2008; Schanzenbächer et al., 2016). Here we demonstrate a second role for transcription in regulating the strength of the network homeostatic response. Loss of Hlf, Tef, and Dbp results in exaggerated homeostatic changes. However, the levels of network activity at baseline are unchanged. This suggests that these transcription factors are activated to restrain homeostatic signaling rather than downregulating network dynamics directly. While a few other genes have been described that have limited effects on network dynamics, but are required for homeostatic plasticity (Shank3, Tatavarty et al., 2020; FMRP, Soden and Chen, 2010; Caspr2, Fernandes et al., 2019), this is, to our knowledge, the first negative regulator of homeostasis to be described.

Present data implicate mEPSC frequency, but not amplitude, as the site of the restraint of homeostatic plasticity. Although two days of activity blockade in dissociated cortical cultures were initially found to produce scaling up of EPSC amplitudes without accompanying changes in mEPSC frequency (Turrigiano et al., 1998), multiple subsequent studies have found evidence for changes in both amplitude and frequency as found here (Echegoyen et al., 2007; Koch et al., 2010; Thiagarajan et al., 2005; Wierenga et al., 2006). Changes in mini frequency are typically interpreted as reflecting increased presynaptic release, either via increases in the numbers of synapses or active release sites, or via changes in the rate at which spontaneous, action potential independent release occurs at each site. Although the increase observed in mEPSC frequency (1.7-fold greater in TKO than WT), was lower than the relative increase in activity levels, measured from the frequency of calcium transients (2.5-fold greater in TKO than WT at 5 days), these two measures need not be linearly related. First, mini frequency can be distinct from action potential evoked release (Crawford et al., 2017), and second, in a highly recurrent network, the relationship between firing and synaptic strength can be nonlinear (Toyoizumi and Abbott, 2011). We cannot eliminate the contribution of inhibitory neurons as a contributing factor to homeostatic plasticity restraint, since Hlf and Tef are expressed in this neuronal type and are also upregulated by inactivity. Interneuron function may also be regulated by Hlf and Tef and this role may predispose the network to seizures. Dissecting the contribution of different cell types during different developmental windows will require generation of a conditional knock out and selective cell-type specific disruption of individual transcription factors at different times in development. mIPSCs received by excitatory neurons cannot explain the exaggerated network hyperexcitability caused by TTX incubation in TKO slices since neither the amplitude nor frequency of mIPSCs is different in the TTX condition. We note, however, that the amplitude of mIPSCs is decreased at baseline in the untreated cultures. Such decrease of the strength of inhibitory currents is insufficient on its own to induce network excitability, since baseline network activity was unchanged in the TKO, but it may make the network more vulnerable to future insults.

It is tempting to imagine a direct relationship between the exaggerated homeostatic plasticity in TKO cortical slices and the increased seizures and spontaneous death of TKO animals. The lack of constraint of homeostatic response can potentially explain, at least in part, the seizure phenotype in animals lacking all three TFs. However, partial genetic compensation of DBP for the survival phenotype but not the homeostatic plasticity phenotype as assayed by calcium imaging, suggests that the propensity to develop seizures and diminished survival have a more complex origin involving other brain regions and perhaps other organs of the body. Gachon et al. (Gachon et al., 2004) first reported that deletion of the PARbZIP proteins TEF, HLF and DBP caused spontaneous and audiogenic seizures. They suggested that this may reflect effects on neurotransmitter metabolism caused by a reduction in pyridoxal phosphate (PLP) since pyridoxal kinase, which catalyzes the last step in the conversion of vitamin B6 into PLP is a target of the PARbZIP proteins. PLP is a required cofactor for decarboxylases and other enzymes involved in the synthesis and metabolism of GABA, glutamate and the biogenic amine neurotransmitters including dopamine, serotonin and histamine. Using HPLC they found no changes in GABA or glutamate, but reduced levels of dopamine, serotonin and histamine in the knockout animals. Although changes in amine transmitters could potentially contribute to phenotypes in vivo, they are unlikely to contribute to plasticity in cortical slice cultures which do not include the brainstem nuclei from which these projections arise. Reduced glutamate decarboxylase action could potentially have contributed to the observed baseline reduction in mIPSCs since heterozygous (Lazarus et al., 2015) and homozygous (Lau and Murthy, 2012) mutants of Gad1, the gene encoding the GAD67 isoform of glutamic acid decarboxylase, have reduced mIPSC amplitudes. However, this does not account for the enhanced homeostatic overshoot following activity blockade, since in the TKO animals, activity blockade did not produce any further reduction in inhibition, but produced robust changes at excitatory synapses.

HLFandTEFform homo- or heterodimersand recognize similar DNA sequences (Gachon, 2007). Here we demonstrate that these TFs can powerfully compensate for the loss of other family members. We varied genotype for all three PARbZip factors and used multiple regression models to show that the peak frequency of calcium transients depends roughly equally on Hlf and Tef but does not appear to depend on Dbp. It also appears that a single allele of either Hlf or Tef in the complete absence of the other factors is sufficient to preserve the normal restraint of homeostatic plasticity, producing a response to activity deprivation that is not significantly different from WT. However, given the large number of genotypes to be tested, the limited number of animals tested may have led us to miss subtle difference that vary with the number of additional alleles, since activity in the presence of 2 alleles was significantly different (albeit with a low effect size) from that in WT. We also found, in agreement with prior findings (Gachon et al., 2004), that Tef and Hlf can robustly compensate for loss of each other and completely restore life span. Such robust compensation and functional redundancy of these TFs highlights their importance and may explain why they have not been identified in GWAS studies as risk factors for epilepsy or other neuropsychiatric diseases.

The mechanisms linking activity deprivation to Tef and Hlf activation were not directly addressed in this work, but seem likely to depend on reduced calcium influx. Prior studies have suggested alternately that scaling up of mEPSC amplitudes depends upon calcium influx through either T-type (Schaukowitch et al., 2017) or L-type (Li et al., 2020) calcium channels. The former pathway has also been found to depend on the activity-dependent transcription factors ELK1 and SRF. Activation of the constraining TFs Tef and Hlf could be secondary or delayed responses dependent on changes in the transcription and translation of other activity-dependent TFs, or may represent a primary response to activity deprivation, analogous to, but in the opposite direction of the well-studied immediate early genes such as Fos, Arc and others.

Besides their function in early embryonic development (Gavriouchkina et al., 2010; Wahlestedt et al., 2017), the PARbZip TFs have been extensively studied for their roles in circadian rhythms in flies, zebrafish, and rodents (Cyran et al., 2003; Vatine et al., 2009; Weger et al., 2021). The present finding that Hlf and Tef also act to restrain homeostatic plasticity is consistent with a recent report demonstrating that PARbZIP TFs are differentially expressed in human epileptogenic tissue (Rambousek et al., 2020). In addition, CLOCK, an upstream regulator of these TFs, is necessary in cortical excitatory neurons for maintaining normal network activity and leads to epilepsy when conditionally deleted in these neurons (Li et al., 2017). Multiple components of the molecular clock are robustly expressed in the neocortex (Bering et al., 2018; Gachon et al., 2004; Kobayashi et al., 2015), consistent with the idea that they might have an additional function in the cortex that is distinct from their circadian role in SCN. However, the role of core clock genes in homeostatic plasticity has not yet been explored. In this work we expand the role of the PARbZIP family of transcription factors to include the negative regulation of homeostatic plasticity.

Why should homeostatic plasticity be subject to such dual “push/pull” regulation? Perhaps because the changes in drive and excitability which must be buffered vary so widely during development. During the first few weeks of postnatal development in rodents, cortical neurons go from receiving few synapses to receiving thousands. This may require developmental downregulation of the strength of homeostatic plasticity. Neuropsychiatric diseases, such as monogenic causes of ASD, can trigger homeostatic changes in circuit properties which restore overall firing rates, but nonetheless have maladaptive effects on cortical function and flexibility (Antoine et al., 2019; Nelson and Valakh, 2015). Conversely, loss of function of genes giving rise to developmental disorders can cause failures of homeostatic plasticity (Blackman et al., 2012; Genç et al., 2020). Our findings provide a potential target for enhancing homeostatic plasticity in contexts where it is insufficient, or for downregulating it when it is maladaptive, and therefore provide an avenue for identifying how the positive and negative regulators of homeostasis interact to stabilize network activity.

## Methods and Materials

### Animals

All procedures were approved by the Institutional Animal Care and Use Committee at Brandeis University (Protocol 20002), and conformed to the National Institutes of Health Guide for the Care and Use of Laboratory Animals. The initial RNAseq screen (Figure 1) was performed using the Cre-dependent tdTomato reporter strain Ai9 (Madisen et al., 2010) crossed with either parvalbumin-ires-Cre (Hippenmeyer et al., 2005), or Emx1-ires-Cre animals (Gorski et al., 2002) obtained from Jackson Labs. All other experiments were performed using wild-type (WT; C57BL/6J) and previously published triple knockout (TKO) Hlf-/-/Dbp-/-/Tef-/- animals (Gachon et al., 2004) which were acquired from the European Mouse Mutant Archive (stock EM:02489) as frozen embryosand propagated via IVF using a provider-recommended protocol. TKO mice were housed on a 12/12 light/dark cycle in a dedicated, climate-controlled facility. Cages were enriched with huts, chew sticks, and tubes. Food and water were available ad libitum, and animals were housed in groups of 2-4 after weaning at p21. Mice of both sexes were used for experiments.

### Organotypic Slice Culture

Organotypic slices were dissected from P6-8 pups. Animals were anaesthetized with a ketamine (20 mg/mL), xylazine (2.5 mg/mL) and acepromazine (0.5 mg/mL) mixture (40 ***μ***Liter, via intraperitoneal injection), the brain was extracted, embedded in 2% agarose, and coronal slices containing primary somatosensory cortex were cut on a compresstome (Precisionary Instruments, Greenville, NC)to 300 μm in ice-chilled ACSF(126mM NaCl,25mM NaHCO3,3mM KCl, 1 mM NaH2PO4, 25 mM dextrose, 2 mM CaCl2 and 2 mM MgCl2, 315-319 mOsm) with a ceramic blade and placed directly onto 6-well Millipore Millicell cell culture inserts (Millipore Sigma PICM0RG50, Burlington, Massachusetts) over 1 mL warmed neuronal media (1x MEM (Millipore-Sigma), 1x GLUTAMAX (Gibco Thermo-Fisher Scientific), 1 mM CaCl2, 2 mM MgSO4,12.9 mM dextrose, 0.08% ascorbic acid, 18 mM NaHCO3, 35 mM 1M HEPES (pH 7.5), 1 μg/mL insulin and 20% Horse Serum (heat inactivated, Millipore-Sigma, Burlington, Massachusetts), pH 7.45 and 305 mOsm). Slices were placed in media containing 1x PenStrep (Gibco Thermo-Fisher Scientific) and 50 μg/mL gentamicin (Millipore-Sigma) for 24 hours and subsequent media changes were antibiotic-free. The slices were then grown at 35oC and 5% CO2. Media was changed every other day to 1 mL fresh media. For TTX treatment, media containing 500 nM TTX was added during the media change at EP 12. For TTX washout experiments, two additional media changes with TTX-free media were performed 5 minutes apart.

### RNA-sequencing

Cell sorting: Slice cultures were converted into a single cell suspension as previously described (Sugino et al., 2006), with some modifications. Organotypic slice cultures (n=3 from 3 separate animals per condition) were placed in ice cold, oxygenated ACSF with 1% FBS and 5% Trehalose (Saxena et al., 2012), that had been 0.4*μ* filtered, containing blockers (APV, DNQX and TTX) to prevent excitotoxicity, and gently removed from the membrane. The cortex was micro-dissected under a Leica MZ16-F fluorescent microscope and the tissue placed in an oxygenated room temperature bath for 45 minutes, supplemented with 1 mg/ml type XIV protease (Sigma-Aldrich). Afterwards the tissue was moved back to the ACSF solution (without protease) and triturated with fire polished Pasteur pipettes of successfully smaller diameters (600, 300 and 150 *μ*). Samples were then sorted with a BD FACSAria Flow Cytometer. All isolated material for mRNA sequencing was harvested using the picopure RNA isolation kit (Life Technologies) and subjected to an on column DNAase digestion. mRNA libraries were prepared as previously described (O’Toole et al., 2017): amplifying with the Ovation RNA-seq system (Nugen), sonicating with a Covaris S 220 Shearing Device, constructing the libraries with the Ovation Rapid DR multiplex System (Nugen), and quantifying library concentration usingthe Illumina library quantification kit (KAPA biosystems). Samples were sequenced on either the Illumina Nextseq or Hiseq machines to a depth of 25 million reads. Illumina sequencing adapters were trimmed from reads using cutadapt followed by mapping to the mm10 genome with STAR using the ENCODE Long RNA-Seq pipeline’s parameters. Reads mapping to exons of known genes were quantified using featureCounts in the Rsubread package (Liao et al., 2019) and differential expression analysis was conducted using DESeq2 (Love et al., 2014). One of the three Emx1-cre TTX replicates was severely contaminated (high levels of nonneuronal genes and low enrichment of neuronal genes) and was excluded. Adjusted p-values for each gene are reported from a Wald Test evaluating the significance of the coefficient representing the treatment group (TTX or Control) followed by the Benjamini-Hochberg correction.

### Quantitative real-time PCR

RNA was extracted from neocortical regions of slice cultures using Trizol reagent followed by RNA Clean & Concentrator kit (Zymo research R1014). cDNA was synthesized with 0.5 μg of RNA using the cDNA Synthesis Kit (Bio-Rad 170-8891) using random hexamers to generate cDNA from total RNA. Quantitative real-time PCR was performed using Corbett Research RG-6000 Real Time PCR Thermocycler with SYBR Green Supermix (Bio-Rad). The following primer sequences were used: RPL10 (forward) 5’-CACGGCAGAAACGAGACTTT-3’, RPL10 (reverse) 5’-CACGGACGATCCTATTGTCA-3’, GAPDH (forward) 5’-TCAATGAAGGGGTCGTTGAT-3’ GAPDH (reverse) 5’-CGTCCCGTAGACAAAATGGT-3’, HLF (forward) 5’-CGGTCATGGATCTCAGCAG-3’, HLF (reverse) 5’-GTACCTGGATGGTGTCAGGG-3’, TEF (forward) 5’-GAGCATTCTTTGCCTTGGTC-3’, TEF (reverse) 5’-GGATGGTCTTGTCCCAGATG −3’.

### Electrophysiology

Organotypic slice cultures were cut out of the cell culture inserts along with the membrane using a scalpel blade. Slices were transferred to a thermo-regulated recording chamber and continuously perfused with oxygenated ACSF. Neurons were visualized on an Olympus upright epifluorescence microscope with 10x air and 40x water immersion objectives. Visually guided whole cell patchclamp recordings were made using near-infrared differential interference contrast microscopy. Recording pipettes of 3–5 MΩresistance contained internal solution with the following concentrations. For excitatory currents and intrinsic excitability: (in mM) 20 KCl, 100 K-gluconate, 10 HEPES, 4 Mg-ATP, 0.3 Na-GTP, 10 Na-phosphocreatine, and 0.1% biocytin. For inhibitory currents: (in mM) 120 KCl, 10 HEPES, 4 Mg-ATP, 0.3 Na-GTP, 10 Na-phosphocreatine, and 0.1% biocytin. Recordings were collected using an AxoPatch 200B amplifier (Axon Instruments, Foster City, CA), filtered at 10 kHz and were not corrected for liquid junction potentials. Data were collected on a Dell computer using custom software running on Igor Pro (WaveMetrics, Lake Oswego, OR).

#### Intrinsic excitability

Cells were selected at random from layer 5 in primary somatosensory cortex. Whole cell recordings were performed with a K-Gluconate-based internal recording solution. Synaptic currents were blocked by adding picrotoxin (PTX) at 15 μM, 6,7-dinitroquinoxaline-2,3-dione(DNQX) 15 μM, and (2R)-amino-5-phosphonovaleric acid (APV) at 35 μM to standard ACSF to block *γ*-aminobutyric acid (GABA), *α*-amino-3-hydroxy-5-methyl-4-isoxazolepropionic acid (AMPA), and N-methyl-d-aspartate (NMDA) receptors, respectively. Cells were held at −65mV by injecting a small current. 500 ms current injections ranging from −25 to 275 pA in 25pA steps increments at random intensity were delivered every 10 seconds. Adaptation ratio was calculated using custom-written IGOR script as previously described (Hattox and Nelson, 2007). Briefly, we calculated the ratio between the third and last inter-spike interval at two times the threshold current.

#### Synaptic currents

To isolated miniature excitatory AMPA-mediated currents 15 μM PTX, 35 μM APV and 500 nM TTX were added to the extracellular recording solution. To isolate inhibitory currents, 10 μM DNQX, 35 μM APV and 500 nM TTX were added to the extracellular solution. The slices were incubated in TTX only during active recording, which lasted no more than 30 minutes for each slice. Cells were clamped at −65mV with a series resistance of below 20MΩ with no compensation for the liquid junction potential. 10 10-second traces were acquired for each cell. Threshold-based detection of mPSCs and action potentials were implemented using custom-written IGOR scripts equipped with routines for baseline subtraction, custom filtering and measurements of intervals, amplitudes and kinetic properties.

### Calcium imaging

To visualized calcium dynamics, 24 hours after slice preparation, 1 *μ*L of pAAV.Syn.GCaMP6f. WPRE. SV40 (AAV9 capsid serotype, Addgene 100837) was applied to the somatosensory cortex in each hemisphere. At day EP14 slices were cut out along with the membrane and placed into the perfusion chamber mounted on a spinning disc microscope (Leica DMI 6000B (Leica Microsystems, Inc., Buffalo Grove, IL), with an Andor CSU W1 spinning disc unit, using an Andor Neo sCMOS cameras and Andor IQ3 to run the system (Andor Technology PLC, Belfast, N. Ireland). Slices were allowed to acclimate while being perfused with 33 °C oxygenated ACSF with no TTX for 10 minutes. Slices were imaged with a 10x objective centered over the primary somatosensory area. For each slice ten minutes of imaging from a 500 μm square per slice were acquired at 33 frames per second.

#### Calcium Imaging analysis

Cellular somatic ROIs were selected by hand from background using custom MATLAB code (http://github.com/VH-Lab/vhlab-TwoPhoton-matlab) based on a standard deviation projection of the video (Figure 1A, bottom panel). Cells were only considered viable if they: were of a characteristic size and shape for neurons; were bright enough to be clearly distinguished from background; were in focus with clear and sharp edges; and had several bouts of firing within the first minute. Active cells (n=10-100) were selected by the experimenter within our area of imaging. We confirmed by subsampling cells in a subset of WT control experiments with more cells (n=7 slices with 27-43 cells) that for the lowest sample of cells measured (n=10) the estimates of peak frequency and synchrony had a coefficient of variation of the mean (SEM/mean calculated across sub-samples within a slice) of 0.05 and 0.01. From these ROIs, mean intensity per cell at each frame was computed. The raw intensity values were transformed to ΔF/F by dividing by baseline values obtained either where the coefficient of variation within 50 frames was below 0.3 ΔF/F units/frame, or from the 5th percentile of all frames. Since the distribution of event amplitudes across cells was potentially dependent on other experimental factors such as the amount of calcium indicator expressed as well as optical factors such as the depth of the cell in the slice, we decided not to try to infer the magnitude of firing from the amplitude of events, but only periods of elevated firing from the timing of events. To produce a peak frequency measure, noise was reduced by fitting the ΔF/F data with a smoothing spline, and active periods were defined as those with a slope above 0.30 ΔF/F units/frame that were within 30 frames of other frames with positive slope. The number of active periods reflected the number of times each cell was active, though this measure had no ability to predict the number of action potentials that made up each firing period. We excluded cells that failed to fire at any time during our period of recording. In addition, some of our slices exhibited some dimming of signal due to a combination of drift in the Z-dimension and fluorescent photobleaching. Experiments were discarded if this decrement caused firing peaks to fall and remain below the slope threshold in the selected cells. Synchrony was defined as the cross-correlation at delta T = 0 during upstates computed for all pairs of recorded cells and averaged across cells and across upstates.

## Supporting information

Data for Figure 6, Table 1

Data for Figure 2--Figure Supplement 2

Data for Figure 2

Data for Figure 2--Figure Supplement 1

Data for Figure 3

Data for Figure 4

Data for Figure 5

## Data Availability

The RNAseq data have been deposited to the BioSample database Accession Numbers: SAMN31104764, SAMN31104765, SAMN31104766, SAMN31104767, SAMN31104768, SAMN31104769, SAMN31104770, SAMN31104771, SAMN31104772, SAMN31104773, SAMN31104774, SAMN31104775

---

**Figure 2—figure supplement 1.**
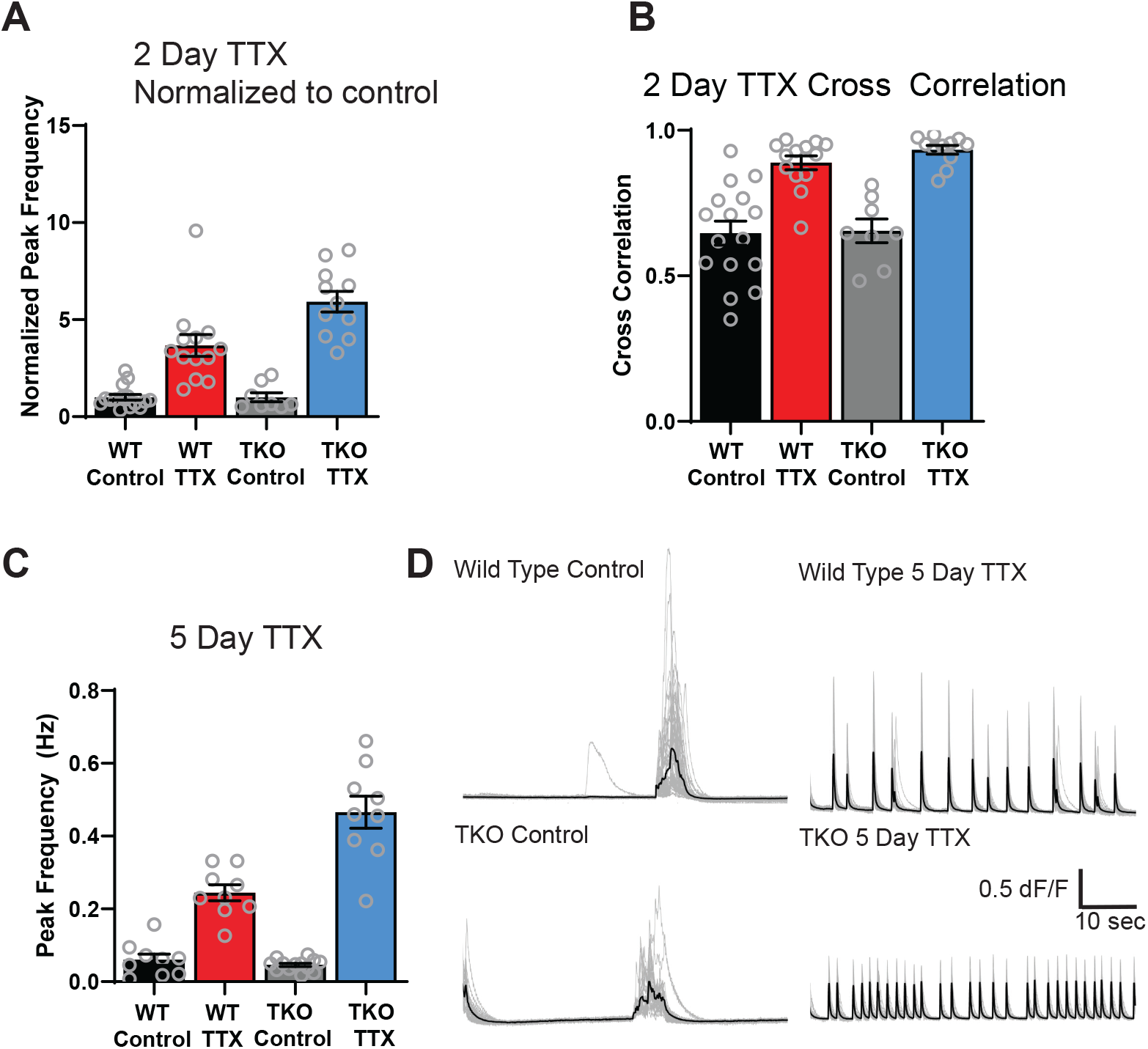
(A) Quantification of peak frequency (from Fig 1D) normalized to control in WT and TKO slices after 2 days of activity deprivation. (B) Quantification of peak synchrony values in each of the conditions in (A). Open symbols are individual slice cultures, filled symbols are mean ± SEM. Synchrony values in WT condition are close to 1 and do not show further increase in the TKO slides. (C) Quantification of peak frequency after 5 days of activity deprivation. and traces from representative slices (D) of the genotypes in (C). Increasing TTX incubation time to 5 days of TTX does not change the TKO phenotype. Peak frequency increased in WT (from 0.06 +/- 0.01 to 0.24 +/- 0.02) and this response is exaggerated in the TKO (from 0.05 +/- 0.005 to 0.47 +/- 0.04;a 2.5-fold greater increase than in WT). As at 2 days, two-way ANOVA revealed significant main effects for TTX (p<0.0001) and genotype (p<0.0001) and a significant interaction between treatment and genotype (p<0.0001) and posthoc test (Tukey) revealed a significant difference between WT TTX and TKO TTX (p<0.0001), but no difference between WT and TKO control activity (p=0.97). Also, as at 2 days, synchrony increased dramatically in the presence of TTX in both genotypes (to 0.96 and 0.93) but this was not significantly different and there was also no significant difference in the values at baseline in the two genotypes (0.54 and 0.58). N=9 for WT control, N=9 for WT TTX, N=13 for TKO control, and N=9 for TKO TTX.

**Figure 2—figure supplement 2.**
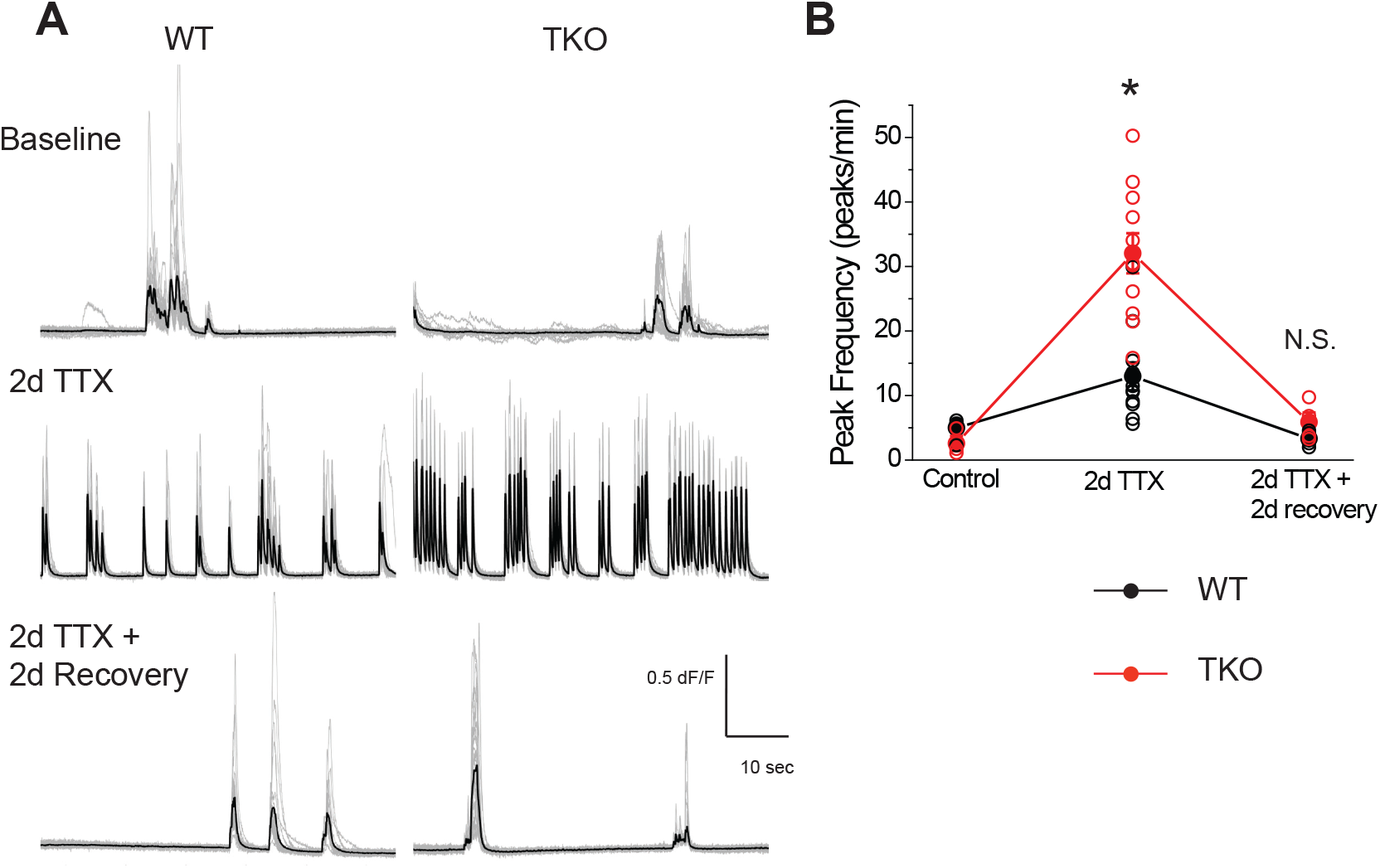
((A) Representative GCaMP6f fluorescence intensity over 1 minute of recording in WT (left panels) and TKO slices (right panels) during control (top row), immediately after TTX washout (middle row) and after two days of recovery following TTX washout (bottom row). (B) Quantification of peak frequency in each of the conditions in (A). Open symbols are individual slice cultures, filled symbols are mean ± SEM. Two-way ANOVA revealed significant main effect for drug treatment (p<0.0001) and genotype (p=0.0009) and a significant interaction between treatment and genotype (p=0.017). Although TTX incubation has a much stronger effect on spontaneous Ca2+ activity immediately after washout in mutant slices (p<0.0001, post-hoc Tukey test), the activity returns to baseline in both genotypes (p=0.99, post-hoc Tukey test). N=5 for WT age-matched control, N=3 for TKO age-matched control, N=11 for WT TTX, N=11 for TKO TTX, N=5 for WT recovery, N=4 for TTX recovery.

## References

Antoine MW, Langberg T, Schnepel P, Feldman DE. Increased Excitation-Inhibition Ratio Stabilizes Synapse and Circuit Excitability in Four Autism Mouse Models. Neuron. 2019 2; 101(4):648–661.e4. doi: 10.1016/j.neuron.2018.12.026, pMID: 30679017.

Bering T, Carstensen MB, Wörtwein G, Weikop P, Rath MF. The Circadian Oscillator of the Cerebral Cortex: Molecular, Biochemical and Behavioral Effects of Deleting the Arntl Clock Gene in Cortical Neurons. Cerebral Cortex. 2018 2; 28(2):644–657. doi: 10.1093/cercor/bhw406, pMID: 28052921.

Blackman MP, Djukic B, Nelson SB, Turrigiano GG. A critical and cell-autonomous role for MeCP2 in synaptic scaling up. The Journal of Neuroscience: The Official Journal of the Society for Neuroscience. 2012 9; 32(39):13529–13536. doi: 10.1523/JNEUROSCI.3077-12.2012, pMID: 23015442 PMCID: PMC3483036.

Crawford DC, Ramirez DMO, Trauterman B, Monteggia LM, Kavalali ET. Selective molecular impairment of spontaneous neurotransmission modulates synaptic efficacy. Nature Communications. 2017 2; 8:14436. doi: 10.1038/ncomms14436, pMID: 28186166 PMCID: PMC5311059.

Cyran SA, Buchsbaum AM, Reddy KL, Lin MC, Glossop NRJ, Hardin PE, Young MW, Storti RV, Blau J. vrille, Pdp1, and dClock form a second feedback loop in the Drosophila circadian clock. Cell. 2003 2; 112(3):329–341. doi: 10.1016/s0092-8674(03)00074-6, pMID: 12581523.

Davis GW. Homeostatic control of neural activity: from phenomenology to molecular design. Annual Review of Neuroscience. 2006; 29:307–323. doi: 10.1146/annurev.neuro.28.061604.135751, pMID: 16776588.

Delvendahl I, Müller M. Homeostatic plasticity-a presynaptic perspective. Current Opinion in Neurobiology. 2019 2; 54:155–162. doi: 10.1016/j.conb.2018.10.003, pMID: 30384022.

Desai NS, Cudmore RH, Nelson SB, Turrigiano GG. Critical periods for experience-dependent synaptic scaling in visual cortex. Nature Neuroscience. 2002 8; 5(8):783–789. doi: 10.1038/nn878, pMID: 12080341.

Desai NS, Rutherford LC, Turrigiano GG. Plasticity in the intrinsic excitability of cortical pyramidal neurons. Nature Neuroscience. 1999 6; 2(6):515–520. doi: 10.1038/9165.

Dörrbaum AR, Alvarez-Castelao B, Nassim-Assir B, Langer JD, Schuman EM. Proteome dynamics during homeostatic scaling in cultured neurons. eLife. 2020 4; 9:e52939. doi: 10.7554/eLife.52939, pMID: 32238265 PMCID: PMC7117909.

Echegoyen J, Neu A, Graber KD, Soltesz I. Homeostatic plasticity studied using in vivo hippocampal activity-blockade: synaptic scaling, intrinsic plasticity and age-dependence. PloS One. 2007 8; 2(8):e700. doi: 10.1371/journal.pone.0000700, pMID: 17684547 PMCID: PMC1933594.

Fernandes D, Santos SD, Coutinho E, Whitt JL, Beltrão N, Rondão T, Leite MI, Buckley C, Lee HK, Carvalho AL. Disrupted AMPA Receptor Function upon Genetic- or Antibody-Mediated Loss of Autism-Associated CASPR2. Cerebral Cortex (New York, NY). 2019 12; 29(12):4919–4931. doi: 10.1093/cercor/bhz032, pMID: 30843029 PMCID: PMC7963114.

Gachon F. Physiological function of PARbZip circadian clock-controlled transcription factors. Annals of Medicine. 2007 1; 39(8):562–571. doi: 10.1080/07853890701491034, pMID: 17852034.

Gachon F, Fonjallaz P, Damiola F, Gos P, Kodama T, Zakany J, Duboule D, Petit B, Tafti M, Schibler U. The loss of circadian PAR bZip transcription factors results in epilepsy. Genes & Development. 2004 6; 18(12):1397–1412. doi: 10.1101/gad.301404, pMID: 15175240 PMCID: PMC423191.

Galvan CD, Hrachovy RA, Smith KL, Swann JW. Blockade of Neuronal Activity During Hippocampal Development Produces a Chronic Focal Epilepsy in the Rat. Journal of Neuroscience. 2000 4; 20(8):2904–2916. doi: 10.1523/JNEUROSCI.20-08-02904.2000, pMID: 10751443.

Gavriouchkina D, Fischer S, Ivacevic T, Stolte J, Benes V, Dekens MPS. Thyrotroph embryonic factor regulates light-induced transcription of repair genes in zebrafish embryonic cells. PloS One. 2010 9; 5(9):e12542. doi: 10.1371/journal.pone.0012542, pMID: 20830285 PMCID: PMC2935359.

Genç, AnJY, Fetter RD, Kulik Y, Zunino G, Sanders SJ, Davis GW. Homeostatic plasticity fails at the intersection of autism-gene mutations and a novel class of common genetic modifiers. eLife. 2020 7; 9:e55775. doi: 10.7554/eLife.55775, pMID: 32609087 PMCID: PMC7394548.

Goold CP, Nicoll RA. Single-cell optogenetic excitation drives homeostatic synaptic depression. Neuron. 2010 11; 68(3):512–528. doi: 10.1016/j.neuron.2010.09.020, pMID: 21040851 PMCID: PMC3111089.

Gorski JA, Talley T, Qiu M, Puelles L, Rubenstein JLR, Jones KR. Cortical Excitatory Neurons and Glia, But Not GABAergic Neurons, Are Produced in the Emx1-Expressing Lineage. The Journal of Neuroscience. 2002 8; 22(15):6309–6314. doi: 10.1523/JNEUROSCI.22-15-06309.2002.

Hattox AM, Nelson SB. Layer V Neurons in Mouse Cortex Projecting to Different Targets Have Distinct Physiological Properties. Journal of Neurophysiology. 2007 12; 98(6):3330–3340. doi: 10.1152/jn.00397.2007, pMID: 17898147.

Hawkins NA, Kearney JA. Confirmation of an epilepsy modifier locus on mouse chromosome 11 and can-didate gene analysis by RNA-Seq. Genes, Brain and Behavior. 2012; 11(4):452–460. doi: 10.1111/j.1601-183X.2012.00790.x, pMID: 22471526 PMCID: PMC3370141.

Hawkins NA, Kearney JA. Hlf is a genetic modifier of epilepsy caused by voltage-gated sodium channel mutations. Epilepsy Research. 2016 1; 119:20–23. doi: 10.1016/j.eplepsyres.2015.11.016.

Hippenmeyer S, Vrieseling E, Sigrist M, Portmann T, Laengle C, Ladle DR, Arber S. A Developmental Switch in the Response of DRG Neurons to ETS Transcription Factor Signaling. PLoS Biology. 2005 5; 3(5):e159. doi: 10.1371/journal.pbio.0030159, pMID: 15836427 PMCID: PMC1084331.

Hobbiss AF, Ramiro-Cortés Y, Israely I. Homeostatic Plasticity Scales Dendritic Spine Volumes and Changes the Threshold and Specificity of Hebbian Plasticity. iScience. 2018 9; 8:161–174. doi: 10.1016/j.isci.2018.09.015, pMID: 30317078 PMCID: PMC6187013.

Ibata K, Sun Q, Turrigiano GG. Rapid Synaptic Scaling Induced by Changes in Postsynaptic Firing. Neuron. 2008 3; 57(6):819–826. doi: 10.1016/j.neuron.2008.02.031, pMID: 18367083.

Johnson HA, Buonomano DV. Development and Plasticity of Spontaneous Activity and Up States in Cortical Organotypic Slices. Journal of Neuroscience. 2007 5; 27(22):5915–5925. doi: 10.1523/JNEUROSCI.0447-07.2007, pMID: 17537962.

Karmarkar UR, Buonomano DV. Different forms of homeostatic plasticity are engaged with distinct tem-poral profiles. The European Journal of Neuroscience. 2006 3; 23(6):1575–1584. doi: 10.1111/j.1460-9568.2006.04692.x, pMID: 16553621.

Kilman V, Rossumv MCW, Turrigiano GG. Activity Deprivation Reduces Miniature IPSC Amplitude by Decreasing the Number of Postsynaptic GABAA Receptors Clustered at Neocortical Synapses. Journal of Neuroscience. 2002 2; 22(4):1328–1337. doi: 10.1523/JNEUROSCI.22-04-01328.2002, pMID: 11850460.

Kim J, Alger BE. Reduction in endocannabinoid tone is a homeostatic mechanism for specific inhibitory synapses. Nature Neuroscience. 2010 5; 13(5):592–600. doi: 10.1038/nn.2517, pMID: 20348918 PMCID: PMC2860695.

Kobayashi Y, Ye Z, Hensch TK. Clock genes control cortical critical period timing. Neuron. 2015 4; 86(1):264–275. doi: 10.1016/j.neuron.2015.02.036, pMID: 25801703 PMCID: PMC4392344.

Koch H, Huh SE, Elsen FP, Carroll MS, Hodge RD, Bedogni F, Turner MS, Hevner RF, Ramirez JM. Prostaglandin E2-Induced Synaptic Plasticity in Neocortical Networks of Organotypic Slice Cultures. Journal of Neuroscience. 2010 9; 30(35):11678–11687. doi: 10.1523/JNEUROSCI.4665-09.2010, pMID: 20810888.

Lambo ME, Turrigiano GG. Synaptic and Intrinsic Homeostatic Mechanisms Cooperate to Increase L2/3 Pyramidal Neuron Excitability during a Late Phase of Critical Period Plasticity. Journal of Neuroscience. 2013 5; 33(20):8810–8819. doi: 10.1523/JNEUROSCI.4502-12.2013, pMID: 23678123.

Lau CG, Murthy VN. Activity-Dependent Regulation of Inhibition via GAD67. Journal of Neuroscience. 2012 6; 32(25):8521–8531. doi: 10.1523/JNEUROSCI.1245-12.2012, pMID: 22723692.

Lazarus MS, Krishnan K, Huang ZJ. GAD67 deficiency in parvalbumin interneurons produces deficits in in-hibitory transmission and network disinhibition in mouse prefrontal cortex. Cerebral Cortex (New York, NY: 1991). 2015 5; 25(5):1290–1296. doi: 10.1093/cercor/bht322, pMID: 24275833 PMCID: PMC4481616.

Li B, Suutari BS, Sun SD, Luo Z, Wei C, Chenouard N, Mandelberg NJ, Zhang G, Wamsley B, Tian G, Sanchez S, You S, Huang L, Neubert TA, Fishell G, Tsien RW. Neuronal Inactivity Co-opts LTP Machinery to Drive Potassium Channel Splicing and Homeostatic Spike Widening. Cell. 2020 6; 181(7):1547–1565.e15. doi: 10.1016/j.cell.2020.05.013, pMID: 32492405 PMCID: PMC9310388.

Li P, Fu X, Smith NA, Ziobro J, Curiel J, Tenga MJ, Martin B, Freedman S, Cea-Del Rio CA, Oboti L, Tsuchida TN, Oluigbo C, Yaun A, Magge SN, O’Neill B, Kao A, Zelleke TG, Depositario-Cabacar DT, Ghimbovschi S, Knoblach S, et al. Loss of CLOCK Results in Dysfunction of Brain Circuits Underlying Focal Epilepsy. Neuron. 2017 10; 96(2):387–401.e6. doi: 10.1016/j.neuron.2017.09.044, pMID: 29024662 PMCID: PMC6233318.

Liao Y, Smyth GK, Shi W. The R package Rsubread is easier, faster, cheaper and better for alignment and quantification of RNA sequencing reads. Nucleic Acids Research. 2019 5; 47(8):e47. doi: 10.1093/nar/gkz114.

Love MI, Huber W, Anders S. Moderated estimation of fold change and dispersion for RNA-seq data with DESeq2. Genome Biology. 2014 12; 15(12):550. doi: 10.1186/s13059-014-0550-8.

Madisen L, Zwingman TA, Sunkin SM, Oh SW, Zariwala HA, Gu H, Ng LL, Palmiter RD, Hawrylycz MJ, Jones AR, Lein ES, Zeng H. A robust and high-throughput Cre reporting and characterization system for the whole mouse brain. Nature Neuroscience. 2010 1; 13(1):133–140. doi: 10.1038/nn.2467, pMID: 20023653 PMCID: PMC2840225.

Mitsui S, Yamaguchi S, Matsuo T, Ishida Y, Okamura H. Antagonistic role of E4BP4 and PAR proteins in the circadian oscillatory mechanism. Genes & Development. 2001 4; 15(8):995–1006. doi: 10.1101/gad.873501, pMID: 11316793.

Moore AR, Richards SE, Kenny K, Royer L, Chan U, Flavahan K, Van Hooser SD, Paradis S. Rem2 stabilizes intrinsic excitability and spontaneous firing in visual circuits. eLife. 2018 5; 7:e33092. doi: 10.7554/eLife.33092, pMID: 29809135 PMCID: PMC6010341.

Nelson SB, Valakh V. Excitatory/Inhibitory Balance and Circuit Homeostasis in Autism Spectrum Disorders. Neuron. 2015 8; 87(4):684–698. doi: 10.1016/j.neuron.2015.07.033, pMID: 26291155 PMCID: PMC4567857.

O’Toole SM, Ferrer MM, Mekonnen J, Zhang H, Shima Y, Ladle DR, Nelson SB. Dicer maintains the identity and function of proprioceptive sensory neurons. Journal of Neurophysiology. 2017 3; 117(3):1057–1069. doi: 10.1152/jn.00763.2016, pMID: 28003412 PMCID: PMC5338617.

Rambousek L, Gschwind T, Lafourcade C, Paterna JC, Dib L, Fritschy JM, Fontana A. Aberrant expression of PAR bZIP transcription factors is associated with epileptogenesis, focus on hepatic leukemia factor. Scientific Reports. 2020 2; 10(1):1–16. doi: 10.1038/s41598-020-60638-7, pMID: 32111960 PMCID: PMC7048777.

Ripperger JA, Shearman LP, Reppert SM, Schibler U. CLOCK, an essential pacemaker component, controls expression of the circadian transcription factor DBP. Genes & Development. 2000 3; 14(6):679–689. doi: 10.1101/gad.14.6.679, pMID: 10733528.

Sanchez-Vives MV, McCormick DA. Cellular and network mechanisms of rhythmic recurrent activity in neocortex. Nature Neuroscience. 2000 10; 3(10):1027–1034. doi: 10.1038/79848, pMID: 11017176.

Saxena A, Wagatsuma A, Noro Y, Kuji T, Asaka-Oba A, Watahiki A, Gurnot C, Fagiolini M, Hensch TK, Carninci P. Trehalose-enhanced isolation of neuronal sub-types from adult mouse brain. BioTechniques. 2012 6; 52(6):381–385. doi: 10.2144/0000113878, pMID: 22668417 PMCID: PMC3696583.

Schanzenbächer C, Sambandan S, Langer J, Schuman E. Nascent Proteome Remodeling following Homeostatic Scaling at Hippocampal Synapses. Neuron. 2016 10; 92(2):358–371. doi: 10.1016/j.neuron.2016.09.058.

Scharfman H. A Novel Animal Model of Epilepsy Caused by Inhibiting Neuronal Activity during Development. Epilepsy Currents. 2002 7; 2(4):127–128. doi: 10.1046/j.1535-7597.2002.t01-1-00048.x, pMID: 15309141 PMCID: PMC321038.

Schaukowitch K, Reese AL, Kim SK, Kilaru G, Joo JY, Kavalali ET, Kim TK. An Intrinsic Transcriptional Pro-gram Underlying Synaptic Scaling during Activity Suppression. Cell Reports. 2017 2; 18(6):1512–1526. doi: 10.1016/j.celrep.2017.01.033, pMID: 28178527.

Soden ME, Chen L. Fragile X Protein FMRP Is Required for Homeostatic Plasticity and Regulation of Synaptic Strength by Retinoic Acid. Journal of Neuroscience. 2010 12; 30(50):16910–16921. doi: 10.1523/JNEUROSCI.3660-10.2010, pMID: 21159962.

Stellwagen D, Malenka RC. Synaptic scaling mediated by glial TNF-*/alpha*. Nature. 2006 4; 440(7087):1054. doi: 10.1038/nature04671.

Stewart M, Silverman JB, Sundaram K, Kollmar R. Causes and Effects Contributing to Sudden Death in Epilepsy and the Rationale for Prevention and Intervention. Frontiers in Neurology. 2020 7; 11:765. doi: 10.3389/fneur.2020.00765, pMID: 32849221 PMCID: PMC7411179.

Styr B, Gonen N, Zarhin D, Ruggiero A, Atsmon R, Gazit N, Braun G, Frere S, Vertkin I, Shapira I, Harel M, Heim LR, Katsenelson M, Rechnitz O, Fadila S, Derdikman D, Rubinstein M, Geiger T, Ruppin E, Slutsky I. Mitochondrial Regulation of the Hippocampal Firing Rate Set Point and Seizure Susceptibility. Neuron. 2019 6; 102(5):1009–1024.e8. doi: 10.1016/j.neuron.2019.03.045, pMID: 31047779.

Sugino K, Hempel CM, Miller MN, Hattox AM, Shapiro P, Wu C, Huang ZJ, Nelson SB. Molecular taxonomy of major neuronal classes in the adult mouse forebrain. Nature Neuroscience. 2006 1; 9(1):99–107. doi: 10.1038/nn1618, pMID: 16369481.

Sun Q, Turrigiano GG. PSD-95 and PSD-93 Play Critical But Distinct Roles in Synaptic Scaling Up and Down. The Journal of Neuroscience. 2011 5; 31(18):6800–6808. doi: 10.1523/JNEUROSCI.5616-10.2011, pMID: 21543610 PMCID: PMC3113607.

Tan HL, Queenan BN, Huganir RL. GRIP1 is required for homeostatic regulation of AMPAR trafficking. Proceedings of the National Academy of Sciences. 2015 8; 112(32):10026–10031. doi: 10.1073/pnas.1512786112, pMID: 26216979.

Tatavarty V, Pacheco AT, Kuhnle CG, Lin H, Koundinya P, Miska NJ, Hengen KB, Wagner FF, Hooser SDV, Turrigiano GG. Autism-associated Shank3 is essential for homeostatic compensation in rodent V1. Neuron. 2020 6; 106(5):769–777.e4. doi: 10.1016/j.neuron.2020.02.033, pMID: 32199104 PMCID: PMC7331792.

Thiagarajan TC, Lindskog M, Tsien RW. Adaptation to synaptic inactivity in hippocampal neurons. Neuron. 2005 9; 47(5):725–737. doi: 10.1016/j.neuron.2005.06.037, pMID: 16129401.

Toyoizumi T, Abbott LF. Beyond the edge of chaos: amplification and temporal integration by recurrent networks in the chaotic regime. Physical Review E, Statistical, Nonlinear, and Soft Matter Physics. 2011 11; 84(5 Pt 1):051908. doi: 10.1103/PhysRevE.84.051908, pMID: 22181445 PMCID: PMC5558624.

Turrigiano GG, Leslie KR, Desai NS, Rutherford LC, Nelson SB. Activity-dependent scaling of quantal amplitude in neocortical neurons. Nature. 1998 2; 391(6670):892–896. doi: 10.1038/36103, pMID: 9495341.

Turrigiano GG, Nelson SB. Homeostatic plasticity in the developing nervous system. Nature Reviews Neuroscience. 2004 2; 5(2):97. doi: 10.1038/nrn1327, pMID: 14735113.

Vatine G, Vallone D, Appelbaum L, Mracek P, Ben-Moshe Z, Lahiri K, Gothilf Y, Foulkes NS. Light Directs Zebrafish period2 Expression via Conserved D and E Boxes. PLoS Biology. 2009 10; 7(10):e1000223. doi: 10.1371/jour-nal.pbio.1000223, pMID: 19859524 PMCID: PMC2759001.

Wahlestedt M, Ladopoulos V, Hidalgo I, Sanchez Castillo M, Hannah R, Säwén P, Wan H, Dudenhöffer-Pfeifer M, Magnusson M, Norddahl GL, Göttgens B, Bryder D. Critical Modulation of H ematopoietic Lineage Fate by Hepatic Leukemia Factor. Cell Reports. 2017 11; 21(8):2251–2263. doi: 10.1016/j.celrep.2017.10.112, pMID: 29166614 PMCID: PMC5714592.

Wallace W, Bear MF. A Morphological Correlate of Synaptic Scaling in Visual Cortex. Journal of Neuroscience. 2004 8; 24(31):6928–6938. doi: 10.1523/JNEUROSCI.1110-04.2004, pMID: 15295028.

Wang Q, Chiu SL, Koropouli E, Hong I, Mitchell S, Easwaran TP, Hamilton NR, Gustina AS, Zhu Q, Ginty DD, Huganir RL, Kolodkin AL. Neuropilin-2/PlexinA3 Receptors Associate with GluA1 and Mediate Sema3F-Dependent Homeostatic Scaling in Cortical Neurons. Neuron. 2017 12; 96(5):1084–1098.e7. doi: 10.1016/j.neuron.2017.10.029.

Wang Q, Maillard M, Schibler U, Burnier M, Gachon F. Cardiac hypertrophy, low blood pressure, and low aldosterone levels in mice devoid of the three circadian PAR bZip transcription factors DBP, HLF, and TEF. American Journal of Physiology-Regulatory, Integrative and Comparative Physiology. 2010 10; 299(4):R1013–R1019. doi: 10.1152/ajpregu.00241.2010, pMID: 20686175.

Weger BD, Gobet C, David FPA, Atger F, Martin E, Phillips NE, Charpagne A, Weger M, Naef F, Gachon F. Systematic analysis of differential rhythmic liver gene expression mediated by the circadian clock and feeding rhythms. Proceedings of the National Academy of Sciences of the United States of America. 2021 1; 118(3):e2015803118. doi: 10.1073/pnas.2015803118, pMID: 33452134 PMCID: PMC7826335.

Wierenga CJ, Walsh MF, Turrigiano GG. Temporal Regulation of the Expression Locus of Homeostatic Plasticity. Journal of Neurophysiology. 2006 10; 96(4):2127–2133. doi: 10.1152/jn.00107.2006.

Wingender E, Schoeps T, Haubrock M, Dönitz J. TFClass: a classification of human transcription factors and their rodent orthologs. Nucleic Acids Research. 2015 1; 43(Database issue):D97–D102. doi: 10.1093/nar/gku1064, pMID: 25361979 PMCID: PMC4383905.

Yoshimura T, Rasband MN. Axon initial segments: diverse and dynamic neuronal compartments. Current Opinion in Neurobiology. 2014 8; 27:96–102. doi: 10.1016/j.conb.2014.03.004.

